# The GLUT4 storage vesicle pool is maintained by intracellular nanovesicle traffic

**DOI:** 10.64898/2026.02.16.706132

**Authors:** Elizabeth Courthold, Gabrielle Larocque, Stephen J. Royle

## Abstract

Glucose transporter 4 (GLUT4) is sequestered intracellularly until the action of insulin causes its redistribution to the cell surface. How cells sequester GLUT4, generating and maintaining GLUT4 storage vesicles (GSVs), is still unclear. Intracellular nanovesicles (INVs) operate on various membrane trafficking pathways and show molecular similarity to GSVs. Here we show that GLUT4 and associated GSV cargo proteins (IRAP/LNPEP, cellugyrin/SYNGR2, sortilin/SORT1, and VAMP2) are trafficked by INVs, which we term GLUT4-flavor INVs. The majority of GLUT4 is carried in GLUT4-flavor INVs which represent a small fraction of the total INV pool. We use a method to capture specific vesicles and test their involvement in the redistribution of GLUT4 in response to insulin. Direct capture of defined GSVs ablated the response, however capture of INVs had a smaller effect. We show that the function of GLUT4-flavor INVs is in the sorting of GSV proteins to supply the insulin-responsive GSV pool and that interfering with INV function by TPD54/TPD52L2 depletion results in mistrafficking of GLUT4 to the plasma membrane. These findings establish GLUT4-flavor INVs as the precursor GSVs that are responsible for intracellular sequestration of GLUT4.

## Introduction

The facilitative glucose transporter GLUT4/SLC2A4 is crucial for maintaining whole body glucose homeostasis in humans. GLUT4 is sequestered inside muscle and fat cells by dynamic membrane trafficking through intracellular compartments, including specialized vesicles termed GLUT4 storage vesicles (GSVs) (Bryant et al., 2002). When insulin is released from the pancreas in response to high blood glucose, it stimulates the redistribution of GLUT4 to the plasma membrane via GSVs, allowing glucose to be transported into the cell, thereby lowering blood glucose to normal levels (Fazakerley et al., 2022). Dysregulation of this process can lead to insulin resistance and the development of type 2 diabetes, which is one of the major health crises facing the global population (Kahn, 1992; Stenbit et al., 1997; van Gerwen et al., 2023).

The intracellular trafficking of GLUT4 has been intensively studied but we do not have a clear picture of how GLUT4 is sequestered into insulin-responsive GSVs (Fazakerley et al., 2022; Gould et al., 2020). Following endocytosis, GLUT4 is sorted at the endosome and trafficked retrogradely to generate the GSVs in a step that likely involves the *trans*-Golgi network (TGN) (Shewan et al., 2003; Kandror and Pilch, 2011; Fazakerley et al., 2009; Antonescu et al., 2008). Additionally, GSVs have been proposed to form directly from the ER-Golgi intermediate compartment in a specialized trafficking step involving CHC22 in humans (Camus et al., 2020; Greig et al., 2024). GLUT4 is primarily expressed in adipocytes and muscle cells, however the pathways that form insulin-responsive GSVs are common to all cells, and heterologous expression of GLUT4 in standard cell lines is sufficient to form GSVs (Morris et al., 2020; Bryant and Gould, 2020). This indicates that core mammalian membrane traffic pathways underpin what otherwise appears to be a specialized membrane trafficking system.

A novel class of membrane trafficking vesicles – intracellular nanovesicles (INVs) – were recently described (Larocque et al., 2020). INVs are characterized by their small size (∼35 nm diameter), lack of protein coat, and the presence of a marker protein TPD54/TPD52L2. They operate on several intracellular trafficking routes and have key functions in cell migration, via recycling α5β1 integrins, and in autophagy, via ATG9A-flavor INVs (Larocque et al., 2021; Fesenko et al., 2025). INVs contain a range of transport machinery such as Rab GTPases, R-SNARES and a variety of cargoes (receptors, transporters), and they primarily move by passive diffusion in the cytoplasm (Larocque et al., 2020; Sittewelle and Royle, 2024). Proteomic and functional analysis indicates that INVs are a large superfamily comprising multiple subtypes or ‘flavors’, with ATG9A-flavor INVs estimated to be ∼20 % of the total INV population (Fesenko et al., 2025).

Intriguingly, several components of GSVs including VAMP2, SCAMPs, Rab10, Rab14, cellugyrin/SYNGR2 are found in the INV proteome (Fesenko et al., 2025; Gould et al., 2020). This suggested to us that INVs may be part of the core trafficking machinery that generates GSVs, or that GSVs may actually be a GLUT4-flavor of INV. Here we test this idea, initially by proteomic comparison of GLUT4 vesicles and INVs, which confirmed the overlap in molecular components. We found that GLUT4 and other known GLUT4 vesicle cargoes are present in INVs, and that the majority of GLUT4 is trafficked in INVs. Functional testing revealed that these GLUT4-flavor INVs are not synonymous with the insulin-responsive GSV pool, instead, they are likely to be the precursor GSVs that mediate intracellular GLUT4 sequestration under basal conditions.

## Results

### Proteomic similarity between GLUT4 vesicles and INVs

We began by determining the GLUT4 vesicle proteome. To do this, we used HeLa cells stably expressing HA-GLUT4-GFP which forms insulin-responsive GLUT4 storage vesicles (GSVs) in the absence of endogenous GLUT4 expression (Morris et al., 2020). Using a vesicle isolation and GFP-Trap approach that was employed previously to determine the INV proteome (Fesenko et al., 2025), we isolated GLUT4-containing vesicles and compared them by mass spectrometry with the same procedure performed on WT HeLa cells. Figure 1A shows the proteins enriched in GLUT4-containing vesicles. Of the 222 proteins, there were key proteins known to be important for GLUT4 trafficking: VAMP2, VAMP3, and VAMP8 which are SNAREs involved in GSV fusion (Zhao et al., 2009); Rab10 and Rab14 (Brewer et al., 2016; Reed et al., 2013; Larance et al., 2005; Brumfield et al., 2021); cellugyrin/SYNGR2 and SCAMPs 1-3 which are found on non-insulin responsive GLUT4 vesicles (Kupriyanova and Kandror, 2000; Kupriyanova et al., 2002; Kioumourtzoglou et al., 2015; Sevilla et al., 1997; Laurie et al., 1993). In addition, core GSV proteins including the Rab GAP TBC1D4 and insulin regulated aminopeptidase (IRAP)/LNPEP were enriched. Furthermore, of the 59 GLUT4 vesicle proteins identified proteomically in adipocytes, 19 were captured in our dataset, 13 of which were in the subset of 23 insulin responsive proteins determined in a previous study (Fazakerley et al., 2015).

**Figure 1.**
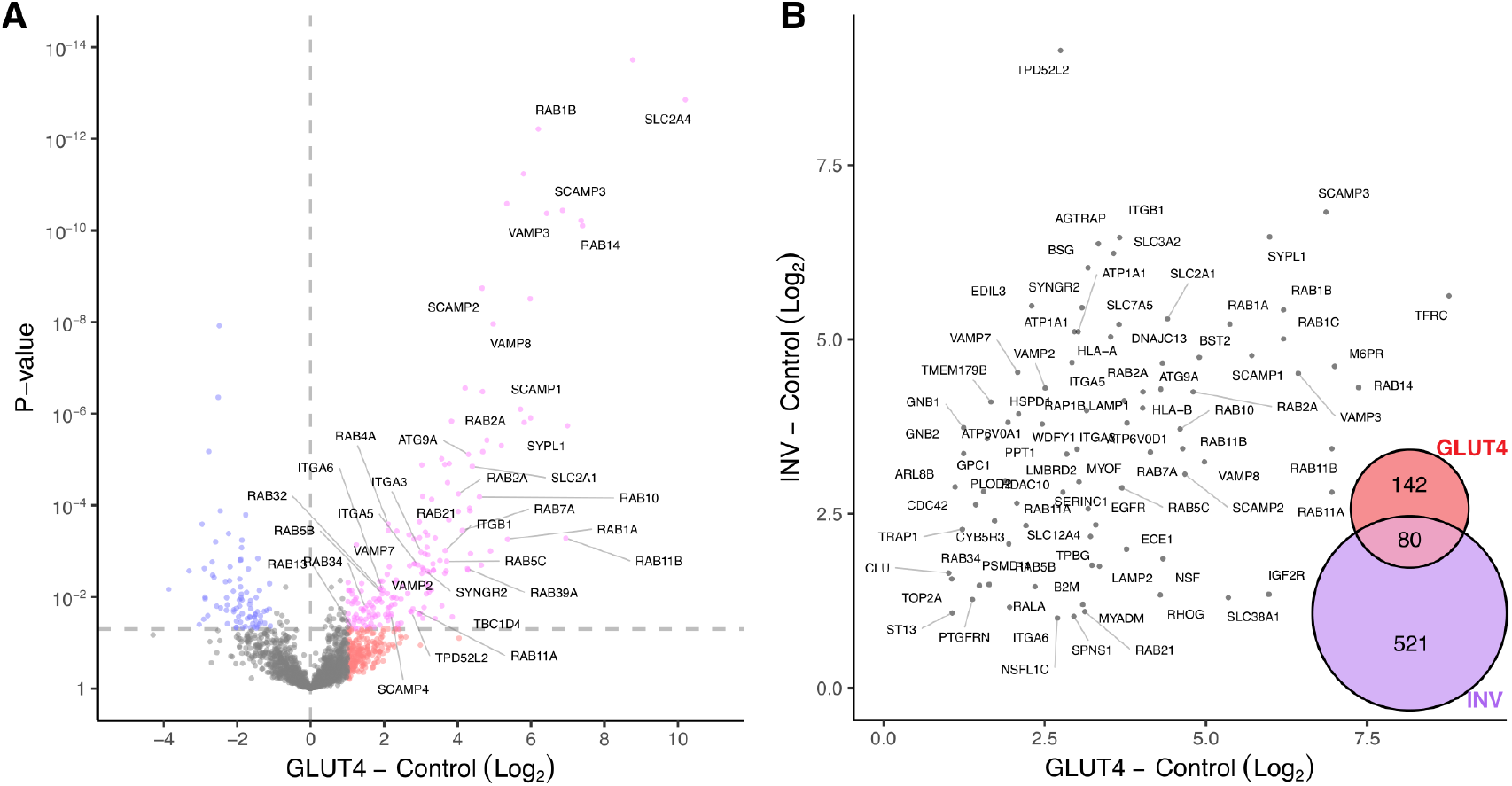
The GLUT4-containing vesicle proteome is similar to the INV proteome. (**A**) Volcano plot to show comparison of proteins detected in GFP-Trap isolation using either HA-GLUT4-GFP HeLa (GLUT4) or HeLa (Control). Data are from three independent runs of three technical replicates each. Colors indicate: pink, fold-change > 2 and *P* < 0.05; red, fold-change > 2 and *P* > 0.05; blue, fold-change < 2 and *P* < 0.05; gray, remainder. For the full list see Supplementary Table S1. (**B**) Comparison of the enrichment of proteins in GLUT4 vs Control, and in INV vs Control. Inset: Euler plot showing the intersection of significantly enriched proteins found in the HA-GLUT4-GFP proteome and the INV proteome, taken from Fesenko et al. (2025).

Importantly, we found the INV marker TPD54/TPD52L2 was significantly enriched (6.7-fold) versus control. Comparison of GLUT4 vesicle and INV proteomes revealed an overlap of 80 proteins which represents 36 % of the total GLUT4 vesicle proteins (Figure 1B). Enrichment was positively correlated (r = 0.36, p = 7.1 × 10^−4^) and included many of the core GSV proteins (Figure 1B). These data suggest that GLUT4 and associated proteins are trafficked via INVs and moreover that GSVs could be a subset of INVs.

### GLUT4 is present in INVs

To directly test for the presence of GLUT4 in INVs, we used a vesicle capture assay described previously (Larocque et al., 2020; Fesenko et al., 2025). Here, rapalog-induced heterodimerization between a vesicle-attached mCherry-FKBP-tagged protein and an FRB-tagged mitochondrial anchor protein (termed MitoTrap), results in the capture of specific vesicles at the mitochondria (Figure 2A). The concomitant co-relocation of a GFP-tagged protein-of-interest, is used to indicate whether that protein is present in the same vesicle. When mCherry-FKBP-TPD54 was relocalized to the mitochondria following rapalog addition, GLUT4-GFP was co-relocated (Figure 2B,C). Co-relocation was not observed when mCherry-FKBP or the mCherry-FKBP-TDP54(R159E) mutant (which does not bind to INVs) was relocalized to the mitochondria (Larocque et al., 2021). This demonstrates that co-relocation of GLUT4 is due to the presence of GLUT4 in INVs captured at the mitochondria, rather than a direct association between GLUT4 and TPD54.

**Figure 2.**
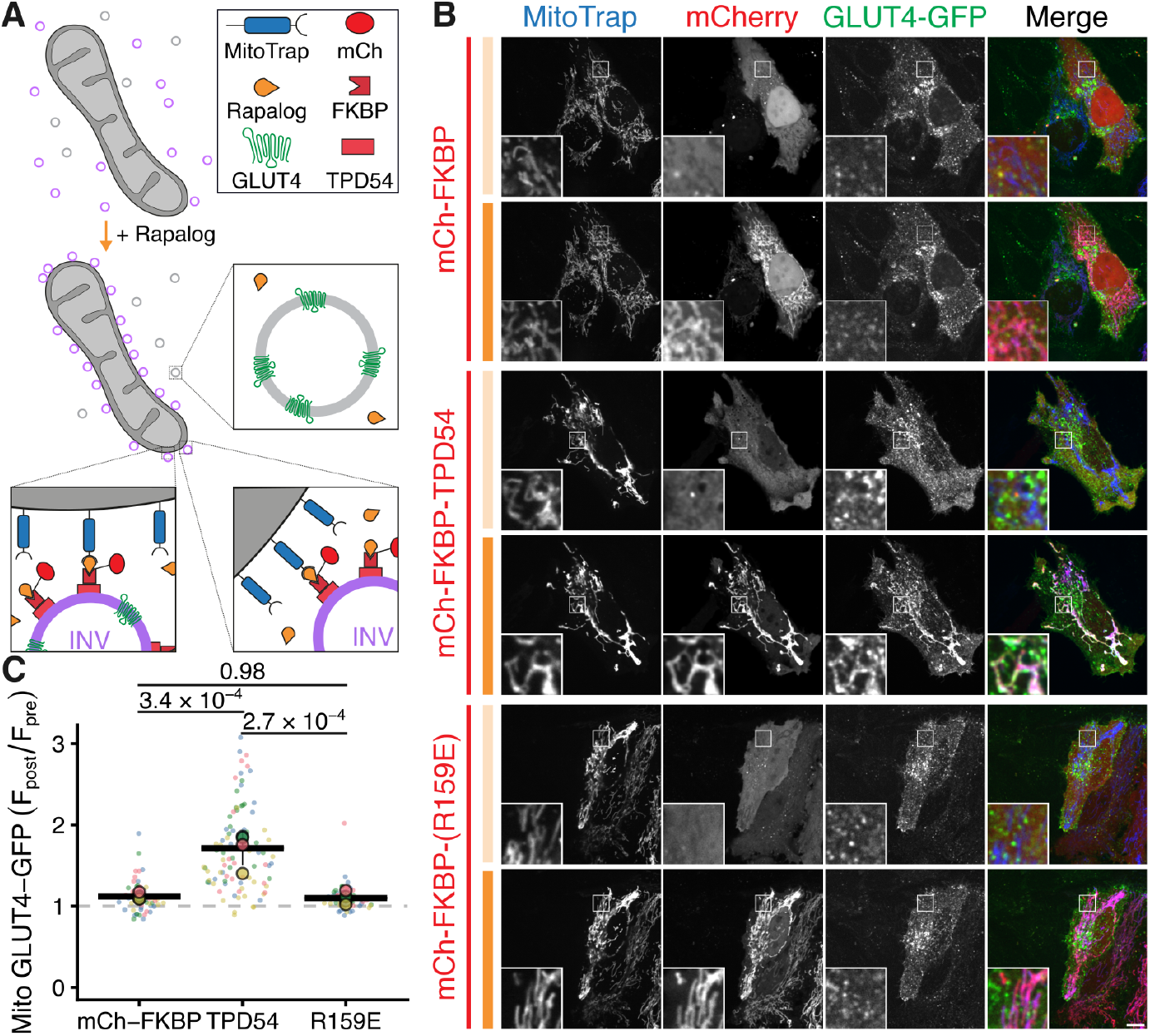
GLUT4 is present in intracellular nanovesicles. (**A**) Schematic diagram of vesicle capture at mitochondria as a test for vesicle co-occupancy. MitoTrap is an FRB domain targeted to mitochondria, mCherry–FKBP–TPD54 is co-expressed and, when rapalog is added, INVs with TPD54 become trapped at the mitochondria. Any protein – GLUT4-GFP in this example – that is also in the INVs will be co-relocated with TPD54 (Larocque et al., 2020). Note that INVs that do not contain GLUT4-GFP are also relocalized, whereas any vesicles that contain GLUT4-GFP but not TPD54, are not captured. (**B**) Representative confocal images of HeLa cells co-expressing GLUT4-GFP (green), MitoTrap (Mito–mCherry–FRB T2098L, blue), and either mCherry–FKBP, mCherry–FKBP–TPD54 WT or mCherry–FKBP–TPD54 R159E (red). Images were taken before and after the addition of Rapalog (5 µM, 10 min). Scale bar, 10 µm; Insets, 4X zoom. (**C**) Superplot to show the change in GLUT4-GFP at the mitochondria after adding rapalog. Points, individual cells; colors, experimental repeat; circles, mean for each repeat; bars, mean ± sd of the experimental means. P-values, ANOVA with Tukey’s *post-hoc* test.

### TPD54 is present on IRAP-positive GLUT4 vesicles

We next performed the reciprocal test – relocalizing GLUT4 vesicles and asking whether TPD54 is co-relocated. Due to poor expression of GLUT4-FKBP-mCherry, we instead used relocalization of mCherry-FKBP-IRAP since IRAP/LNPEP i) co-traffics with GLUT4 (confirmed below), ii) featured in our proteomic data and iii) is commonly used as a marker for GSVs (Garza and Birnbaum, 2000; Ross et al., 1996, 1997; Chen and Lippincott-Schwartz, 2015). When mCherry-FKBP-IRAP was relocalized to the mitochondria following rapalog addition, GFP-TPD54 co-relocated (Figure 3A,B). Co-relocation was similar regardless of whether GFP-TPD54 was endogenously tagged or transiently expressed. In both cases, some cytoplasmic GFP-TPD54 signal remained suggesting that not all INVs carry IRAP, and by extension, GLUT4. No co-relocation was observed for GFP-TPD54(R159E), indicating that co-relocation of GFP-TPD54 was due to the presence of TPD54 on the IRAP-positive GLUT4 vesicles trapped at the mitochondria.

**Figure 3.**
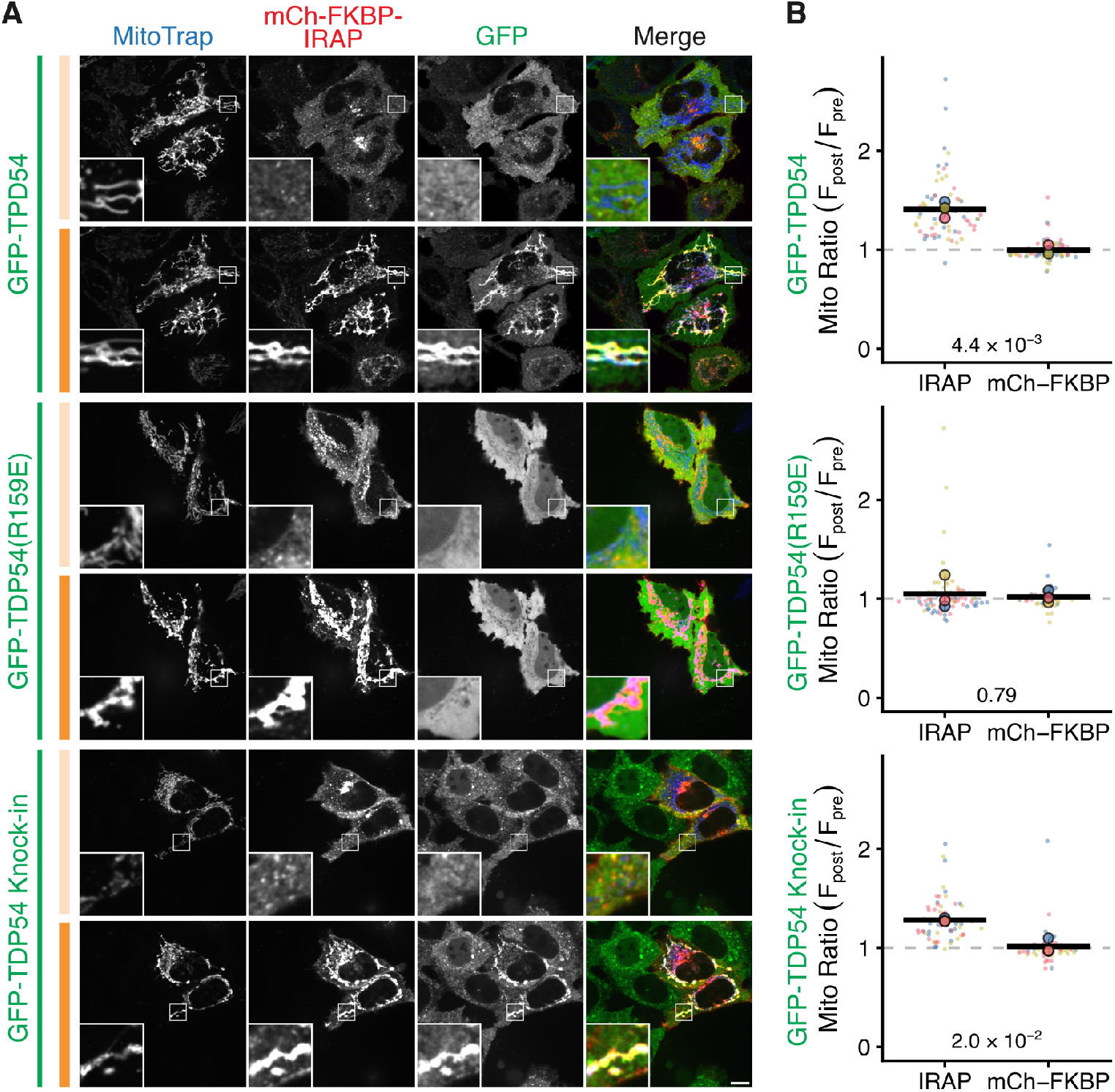
TPD54 is present on IRAP-positive GLUT4 vesicles. (**A**) Representative confocal images of HeLa cells co-expressing MitoTrap (Mito–mCherry–FRB T2098L, blue), mCherry–FKBP–IRAP (red) and either GFP–TPD54, GFP-TPD54(R159E), or GFP-TPD54 knock-in (green). Images were taken before and after the addition of Rapalog (5 µM, 10 min). Insets: 4X zoom. Scale bar, 10 µm. (**B**) Superplots to show the change in fluorescent signal of the GFP-tagged protein at the mitochondria after rapalog addition in mCherry-FKBP-IRAP or mCherry-FKBP (control) conditions. Points, individual cells; colors, experimental repeat; circles, mean for each repeat; bars, mean ± sd of the experimental means. P-values, Students’ t-test with Welch’s correction.

### GLUT4 vesicle cargoes are present in INVs

If GLUT4 vesicles are a subset of INVs then we would expect that other GSV cargoes would be co-relocated when INVs are captured at the mitochondria. First, we confirmed IRAP and GLUT4 can be used interchangeably. We saw that stably expressed HA-GLUT4-GFP was co-relocated by relocalization of mCherry-FKBP-IRAP to the mitochondria, but not by mCherry-FKBP relocalization (Figure 4A,C). Additionally, we observed that IRAP-pHluorin was co-relocated by relocalization of mCherry-FKBP-TPD54 but not the R159E mutants (Figure 4B,C). Second, we used cellugyrin/SYNGR2, VAMP2 and sortilin/SORT1 as exemplar cargoes of GSVs to test for co-relocation. All three co-relocated with mCherry-FKBP-IRAP but not with mCherry-FKBP relocalization, as expected (Figure 4A,C). Similarly, these three cargoes were also co-relocated with mCherry-FKBP-TPD54, but not with TPD54(R159E) (Figure 4B,C). Since these findings indicate that GSV cargoes are present on INVs, they suggest that GSVs may be a subtype of INV.

**Figure 4.**
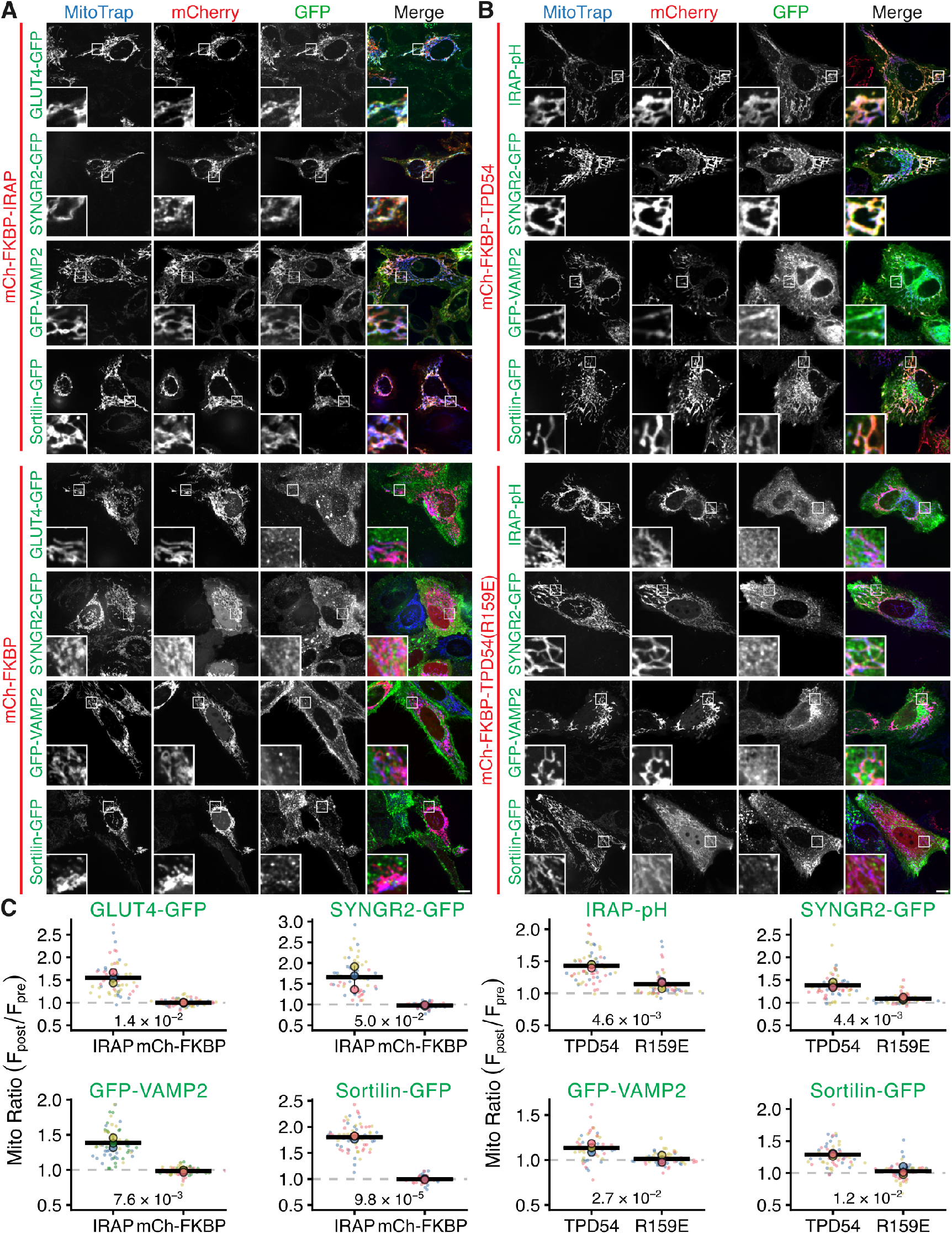
GLUT4 vesicle cargoes are present in intracellular nanovesicles. (**A&B**) Representative confocal images of GFP-tagged GLUT4 vesicle cargoes in HeLa cells co-expressing MitoTrap (Mito–mCherry–FRB(T2098L)) and either mCherry-FKBP-IRAP, mCherry-FKBP (A), mCherry-FKBP-TPD54, or mCherry-FKBP-TPD54(R159E) (B). Images show the cells after the addition of Rapalog (5 µM, 10 min). Scale bar, 10 µm; Insets, 4X zoom. (**C**) Superplot to show the change in GFP-tagged cargo at the mitochondria after adding rapalog. Points, individual cells; colors, experimental repeat; circles, mean for each repeat; bars, mean ± sd of the experimental means. P-values, Students’ t-test with Welch’s correction.

### Vesicle co-occupancy determined by single vesicle tracking

Besides vesicle capture, we wanted an alternative way to designate vesicle subpopulations in living cells. We therefore developed a single vesicle image analysis method that allows us to determine the co-occupancy of proteins in fast-moving subresolution vesicles that are hard to resolve. Briefly, cells expressing two candidate vesicle proteins are imaged at high spatiotemporal resolution with simultaneous two-channel acquisition (Figure 5A). Single vesicles in each channel are tracked independently and then correlated such that we can determine apparent colocalization frame-by-frame with an error rate of 0.05 (Figure 5B,C). From the resultant colocalization histories for each vesicle track, using a criterion of > 80 percent of frames showing colocalization, we determine co-occupancy of the two proteins (Figure 5D); see Methods and Supplementary Figure S1 for more information.

**Figure 5.**
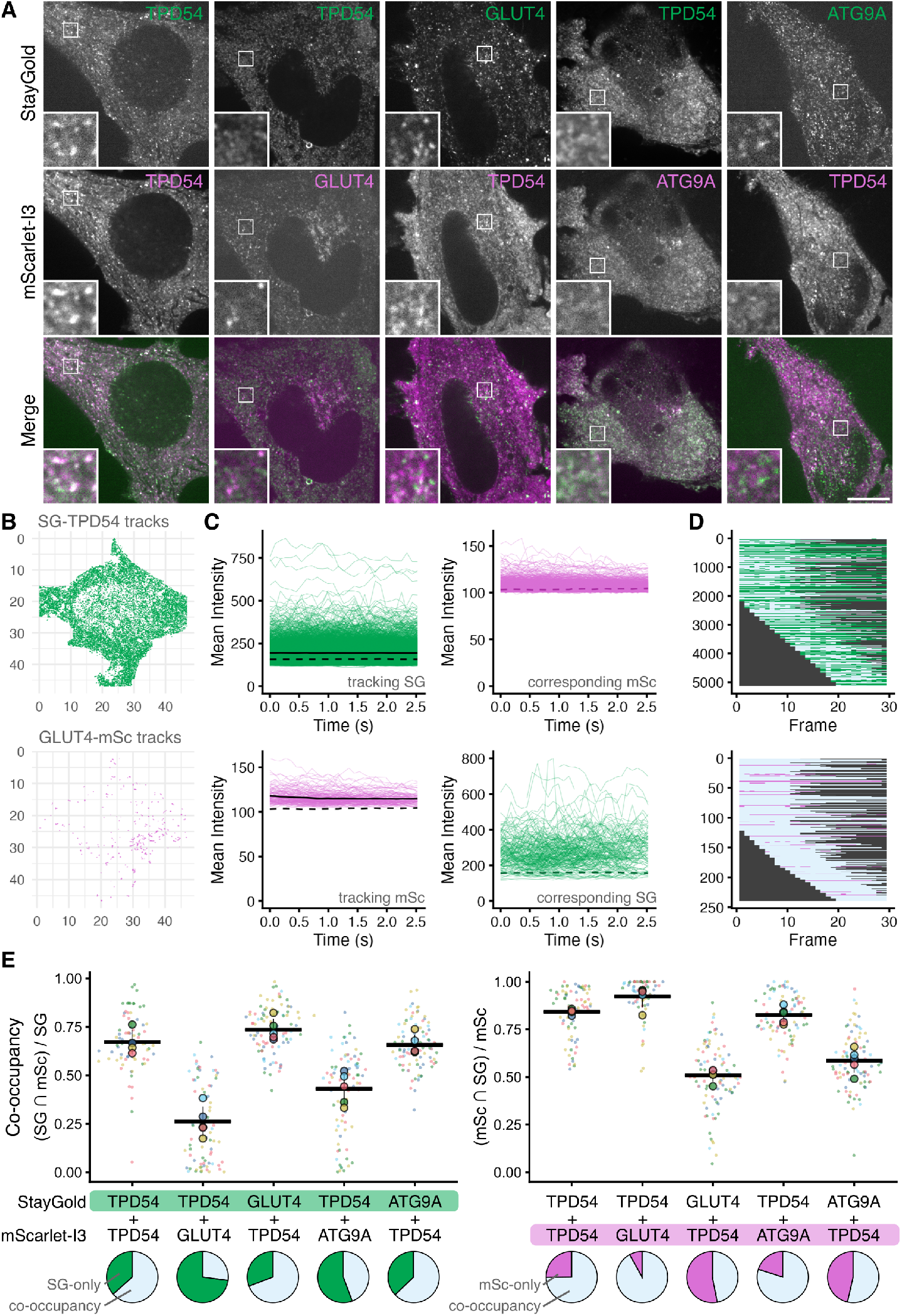
Vesicle co-occupancy determined by single vesicle tracking. (**A**) Stills from typical live-cell dual-color imaging movies. HeLa cells coexpressing the indicated constructs and were imaged at 90 ms per frame. Insets: 4X zoom. Scale bar, 10 µm. See Supplementary Videos SV1, SV2, SV3, SV4, and SV5. (**B**) Example vesicle tracking results for StayGold-TPD54 and GLUT4-mScarlet-I3. Movies were tracked twice, using StayGold (green, above) or mScarlet-I3 (magenta, below) channel. Tracks indicate single vesicle events. (**C**) The fluorescence intensities of the tracked vesicles and corresponding location are shown as a function of time. In the tracked channel, solid and dashed lines indicate the mean and threshold for colocalization (see Methods). The threshold is shown overlaid onto the corresponding data, which determines colocalization. (**D**) Lifetimes of tracked events, blue indicates colocalization while green or magenta, no colocalization. If 80 % of frames show colocalization, then the track is co-occupied. (**E**) Superplots to show the fraction of tracked vesicles that were co-occupied in cells expressing the combination of proteins indicated. Points, individual cells; colors, experimental repeat; circles, mean for each repeat; bars, mean ± sd of the experimental means. Below, pie charts show the aggregate data of tracks that were co-occupied (blue) or not (green or magenta). Total tracks, 9.4 × 10^5^ (SG), 4.8 × 10^5^ (mSc); average tracks per cell, 2261 (SG), 1155 (mSc); n_cell_, 10-22 (per condition per experiment); n_expt_, 5.

As a positive control, we tested StayGold-TPD54 and mScarlet-I3-TPD54 and found that approximately 67– 84 % of green or red tracks were co-occupied (Figure 5E). This range can be taken to represent the maximum co-occupancy we can observe with this method, for a protein combination that is expected to give 100 % co-occupancy in vesicles at extremely high density. Next, to look at GLUT4-INV co-occupancy, we imaged GLUT4/TPD54 pairs, either StayGold-TPD54 with GLUT4-mScarlet-I3 or GLUT4-StayGold with mScarlet-I3-TPD54. The proportion of GLUT4-StayGold or GLUT4-mScarlet-I3 tracks that were co-occupied with TPD54 was 74 % or 92 %; whereas for StayGold-TPD54 or mScarlet-I3-TPD54 tracks, the proportion was only 35 % or 43 %, respectively.

To calibrate the co-occupancy data we also examined ATG9A-INV co-occupancy. To do this we imaged either StayGold-TPD54 with ATG9A-mScarlet-I3 or ATG9A-StayGold with mScarlet-I3-TPD54. As with GLUT4, the fraction of ATG9A-StayGold or ATG9A-mScarlet-I3 tracks that were co-occupied with TPD54 was high, 66 % or 81 %; whereas for StayGold-TPD54 or mScarlet-I3-TPD54 tracks, the proportion was again lower, only 48 % or 52 %, respectively.

These findings support the conclusion that virtually all GLUT4 vesicles are TPD54-positive but that only a subset of INVs carry GLUT4. In our previous work, we estimated that the fraction of INVs that were ATG9A-flavor was approximately 20 % (Fesenko et al., 2025).

Our present analysis indicates the fraction of INVs carrying GLUT4 is smaller than those that carry ATG9A. Together our results indicate that a high proportion of either ATG9A or GLUT4 vesicles are INVs, whereas they each represent a sub-population of INVs.

### Vesicle mobility analysis and direct visualization indicate that cargo-laden INVs are enlarged

Two properties of INVs are their small size and their diffusive motion (Larocque et al., 2020; Sittewelle and Royle, 2024). If GLUT4 vesicles are a subtype of INV, then we would predict that they would be a similar size and that they too would move by free diffusion. We therefore examined single vesicle tracking data to assess these fundamental vesicle properties.

Tracking of StayGold-tagged versions of TPD54, GLUT4 and ATG9A was used to determine the mean diffusion coefficient (*D*) of vesicles per cell, as well as the distribution of *D* for all tracks analyzed, along with the mean squared displacement (MSD) exponent α (Figure 6A-C). Tracks of all three vesicle markers had a similar distribution of α, which centered on 1 each case. This indicates that vesicles with each of the three markers move predominantly by free diffusion, with only small fractions of sub-diffusive and ballistic motions (Figure 6C). However, *D* was more variable. Compared to GLUT4 and ATG9A, TPD54 had a larger mean diffusion coefficient overall, 0.08 µm^2^ s^−1^ (Figure 6A). Note that TPD54 has a broad distribution which encompasses the smaller mean diffusion coefficients of ATG9A and GLUT4 of 0.05 and 0.065 µm^2^ s^−1^, respectively (Figure 6A,B). These results suggests there are faster and slower diffusing INVs, and that INVs carrying ATG9A or GLUT4 are at the slower end of the population.

**Figure 6.**
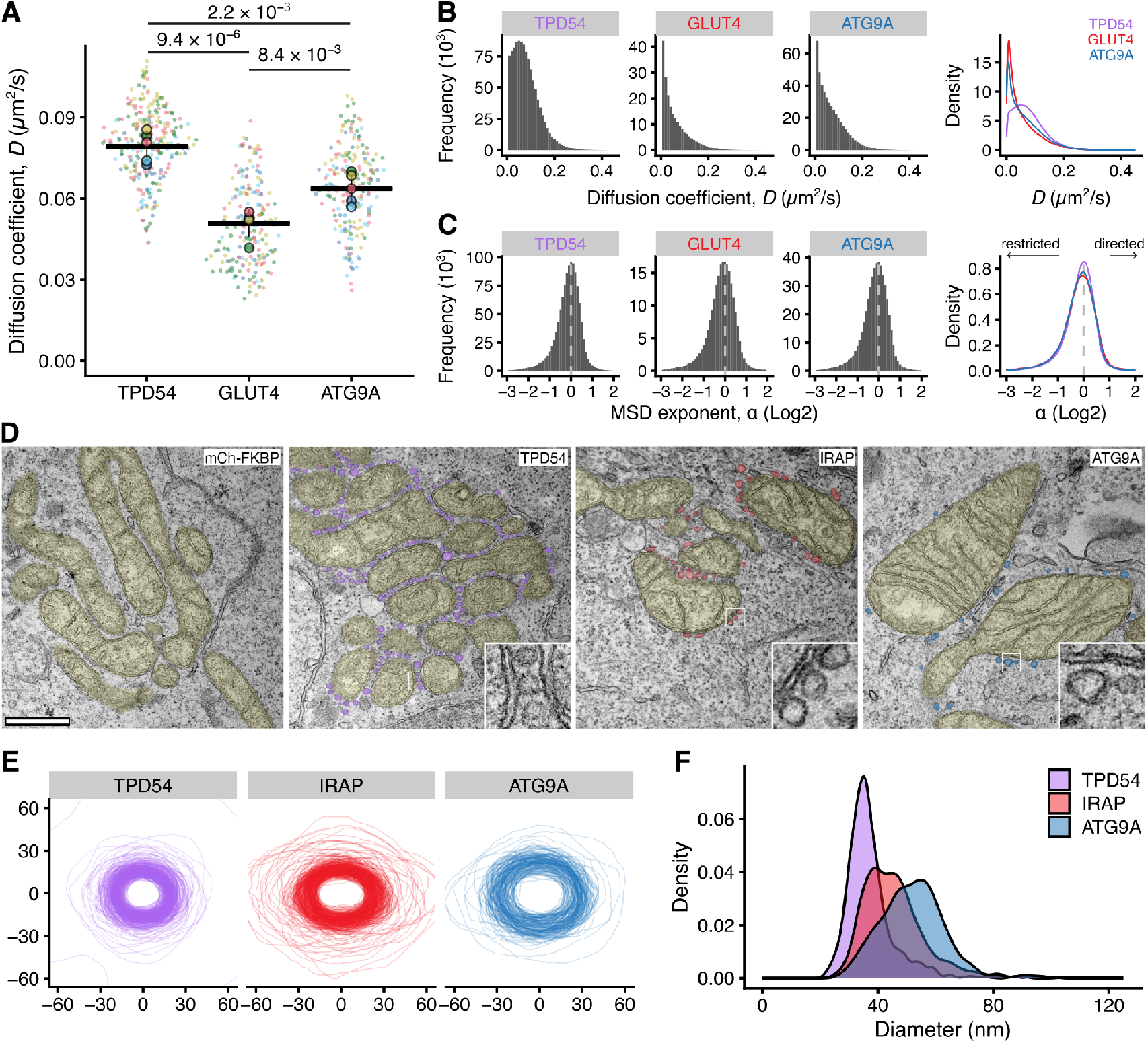
Vesicle mobility analysis and direct visualization indicate that cargo-laden INVs are enlarged. (**A**) Superplot to show the diffusion coefficient (*D*) of single vesicles from tracking of either StayGold-TPD54, GLUT4-StayGold or ATG9A-StayGold in HeLa cells. Points, mean *D* per cell; colors, experimental repeat; circles, average *D* per experiment; bars, mean ±sd of experimental means. P-values, ANOVA with Tukey’s *post-hoc* test. (**B,C**) Histograms and density overlays for all tracks, longer than 0.5 s to show the distribution of *D* (B) and the MSD exponent alpha (*α*) (C). Total tracks, 1.9 × 10^6^; n_cell_, 192-259; n_expt_, 5. (**D**) Representative transmission electron micrographs of cells expressing pMito-GFP-FRB and either mCherry-FKBP (mCh-FKBP, control), mCherry-FKBP-TPD54 (TPD54), ATG9A-FKBP-mCherry (ATG9A) or mCherry-FKBP-IRAP (IRAP) and treated with rapamycin. Mitochondria are false-colored yellow, vesicles attached to mitochondria are purple, red and blue respectively. No vesicles were found attached to mitochondria in mCherry-FKBP. Scale bar, 500 nm. (**E**) Outlines of vesicles attached to mitochondria. A random sample of 150 outlines are shown aligned and rotated with their long axis at *y* = 0. (**F**) Density profiles of all measured vesicles. n_vesicle_ = 390 - 1838.

Since diffusion is inversely proportional to vesicle size (Einstein, 1905; Sittewelle and Royle, 2024), it follows that these differences may reflect a cargo-dependent difference in INV size. To test this notion, we used electron microscopy (EM) to directly visualize vesicles that were relocalized to the mitochondria via mCherry-FKBP-TPD54, mCherry-FKBP-IRAP, or ATG9A-FKBP-mCherry co-expressed with green MitoTrap (Figure 6D). Using a correlative approach, we confirmed by fluorescence microscopy that rapamycin (200 nM) induced the relocalization of each vesicle marker, with mCherry-FKBP being used as a control. The same cells were then visualized by EM allowing us to measure the size of vesicles that were specified according to their cargo (Figure 6D-C). We found that while the full range of mCherry-FKBP-TPD54 vesicle diameters was 21 −180 nm, the median and interquartile range was 35.7 (32.3 – 39.8) nm. This is in agreement with our previous measures of INV size (Larocque et al., 2020). Vesicles captured via mCherry-FKBP-IRAP or ATG9A-FKBP-mCherry were slightly larger on average, 44.2 (38.1 – 50.8) nm or 52.5 (45.4 - 58.6) nm in diameter, respectively; placing them in the upper quartile of the INV size distribution. These results are consistent with our mobility measurements and together they indicate that INVs carrying GLUT4 or ATG9A are larger and diffuse more slowly in the cytosol as a result.

### Insulin decreases the mobility of GLUT4 vesicles

So far, our experiments likely resemble an insulin-stimulated state, due to the presence of serum, which contains insulin. We therefore wanted to examine the effect of insulin treatment on GLUT4 vesicle mobility and on the vesicular co-occupancy of GLUT4 and TPD54. Analysis of GLUT4 vesicle mobility in HeLa cells stably expressing HA-GLUT4-GFP revealed that the diffusion coefficient in basal (serum-starved) conditions was higher when compared with insulin stimulation (Figure 7A,B). Under basal conditions, GLUT4 vesicles had a population mean *D* of approximately 0.1 µm^2^ s^−1^, which is closer to the population mean for INVs (0.08 µm^2^ s^−1^). By contrast, mobility of GLUT4 vesicles during insulin stimulation was slower, 0.05 µm^2^ s^−1^ (Figure 7A). In both cases, α centered on 1 and the distribution of individual track *D* shifted towards slower rates, meaning that the vesicles remained diffusive in either condition and that the slower rate was not due to a larger subdiffusive pool. These data suggest that following insulin stimulation, there is a redistribution of GLUT4 into vesicles which are larger thereby resulting in slower diffusion. We therefore looked at the vesicle co-occupancy of HA-GLUT4-GFP and mScarlet-I3-TPD54 in basal and insulin conditions (Figure 7C). As with the previous GLUT4/TPD54 imaging, the majority of HA-GLUT4-GFP tracks were co-occupied with mScarlet-I3-TPD54 while for mScarlet-I3-TPD54 tracks, the proportion that were co-occupied with HA-GLUT4-GFP was lower. Interestingly, we found that under insulin stimulation, there was a small but significant decrease in co-occupancy of GLUT4 and TPD54 suggesting that GLUT4 moves away from INVs, into a different GLUT4 vesicle population.

**Figure 7.**
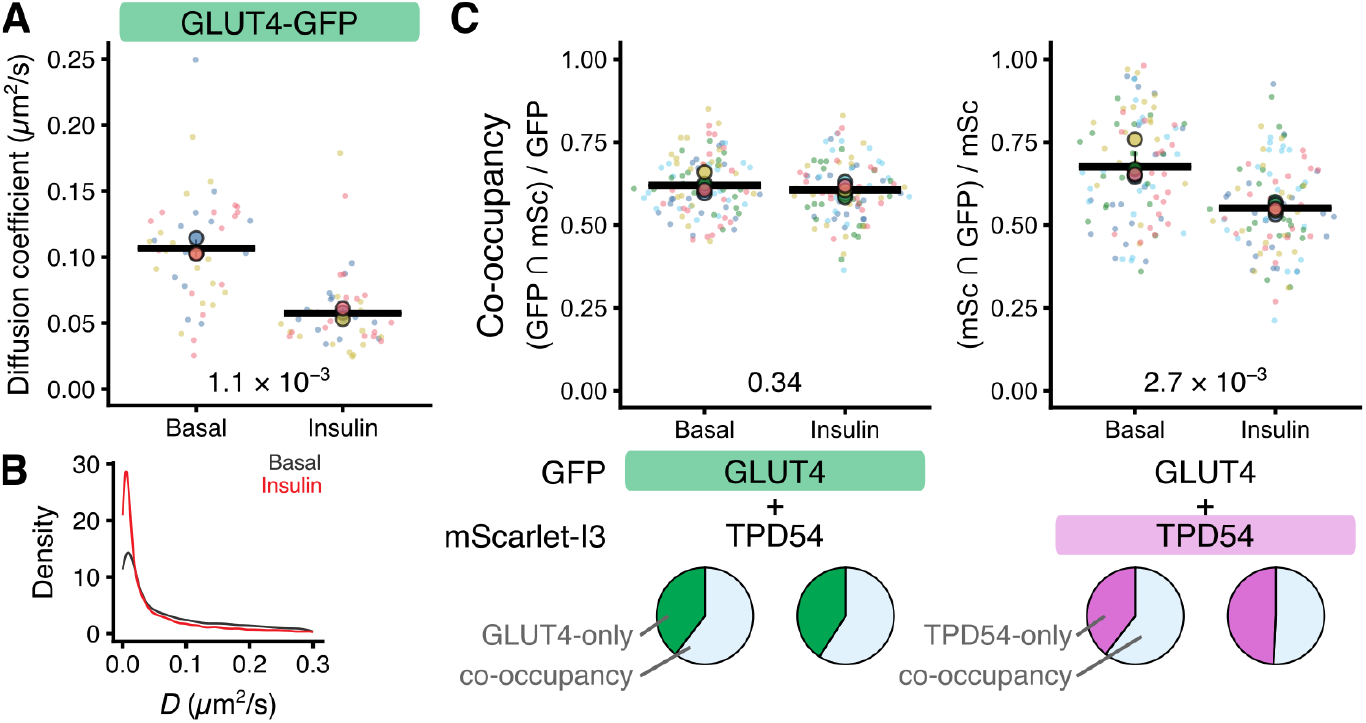
Effect of insulin on the mobility of GLUT4 vesicles and INV-mediated GLUT4 traffic. (**A**) Superplot to show the diffusion coefficient (*D*) of single vesicles from tracking of HeLa cells stably expressing HA-GLUT4-GFP. Cells were either serum-starved (Basal) or treated with insulin (1 µg/mL for 10 min). Points, mean *D* per cell; colors, experimental repeat; circles, average *D* per experiment; bars, mean ± sd of experimental data. P-values, Student’s t-test with Welch’s correction. (**B**) Density overlays for all tracks, longer than 0.5 s to show the distribution of *D*. Total tracks, 2.8 × 10^4^; n_cell_, 43-46; n_expt_, 5. (**C**) Co-occupancy measurement from single vesicle tracking of HeLa cells stably expressing HA-GLUT4-GFP and co-expressed mScarlet-I3-TPD54. Cells were starved or treated with insulin as in A. Superplots to show the fraction of tracked vesicles that were co-occupied in cells expressing the combination of proteins indicated. Points, cells; circles, experimental mean; bars, mean ± sd. P-values, Student’s t-test with Welch’s correction. Below, pie charts show the aggregate data of tracks that were co-occupied (blue) or not (green or magenta). Total tracks, 4.0 × 10^5^ (GFP), 3.2 × 10^5^ (mSc); average tracks per cell, 2028 (GFP), 1571 (mSc); n_cell_, 19-21 (per condition per experiment); n_expt_, 5.

### GLUT4 INVs are distinct from insulin-responsive GLUT4 vesicles

Having determined that GLUT4 and its cargoes are found in INVs and what the properties of those INVs are, we next wanted to determine their function. A key property of GLUT4 is its intracellular retention under basal conditions (absence of insulin), and its rapid redistribution to the cell surface upon insulin stimulation (Bryant et al., 2002). This redistribution can be measured by selective labeling of surface HA-GLUT4-GFP stably expressed in HeLa cells using using immunofluorescence detection of anti-HA applied to non-permeabilized cells, and taking this signal as a ratio of the GFP fluorescence which represents total GLUT4 (Lampson et al., 2000). We reasoned that selective capture of vesicles at the mitochondria could be used to test what effect their confinement has on the insulin-induced redistribution of HA-GLUT4-GFP. As a positive control, we first performed vesicle capture with mCherry-FKBP-IRAP and dark MitoTrap. Cells were starved of insulin and then were treated with rapalog (5 µM, 10 min), followed by 20 min insulin stimulation (1 µg/mL), and then surface and total GLUT4 were measured by immunofluorescence microscopy. In cells not treated with rapalog there was an increase in the fraction of total GLUT4 at the surface following insulin stimulation (Figure 8A,B). In cells where mCherry-FKBP-IRAP-containing vesicles were trapped at the mitochondria with rapalog treatment, this response was substantially reduced. In fact, the surface levels of GLUT4 were lowered under basal conditions by vesicle capture. This likely reflects the maximal effect that 10 min of vesicle confinement can have and confirms that GLUT4 and IRAP co-exist in the insulin-responsive GSV pool.

**Figure 8.**
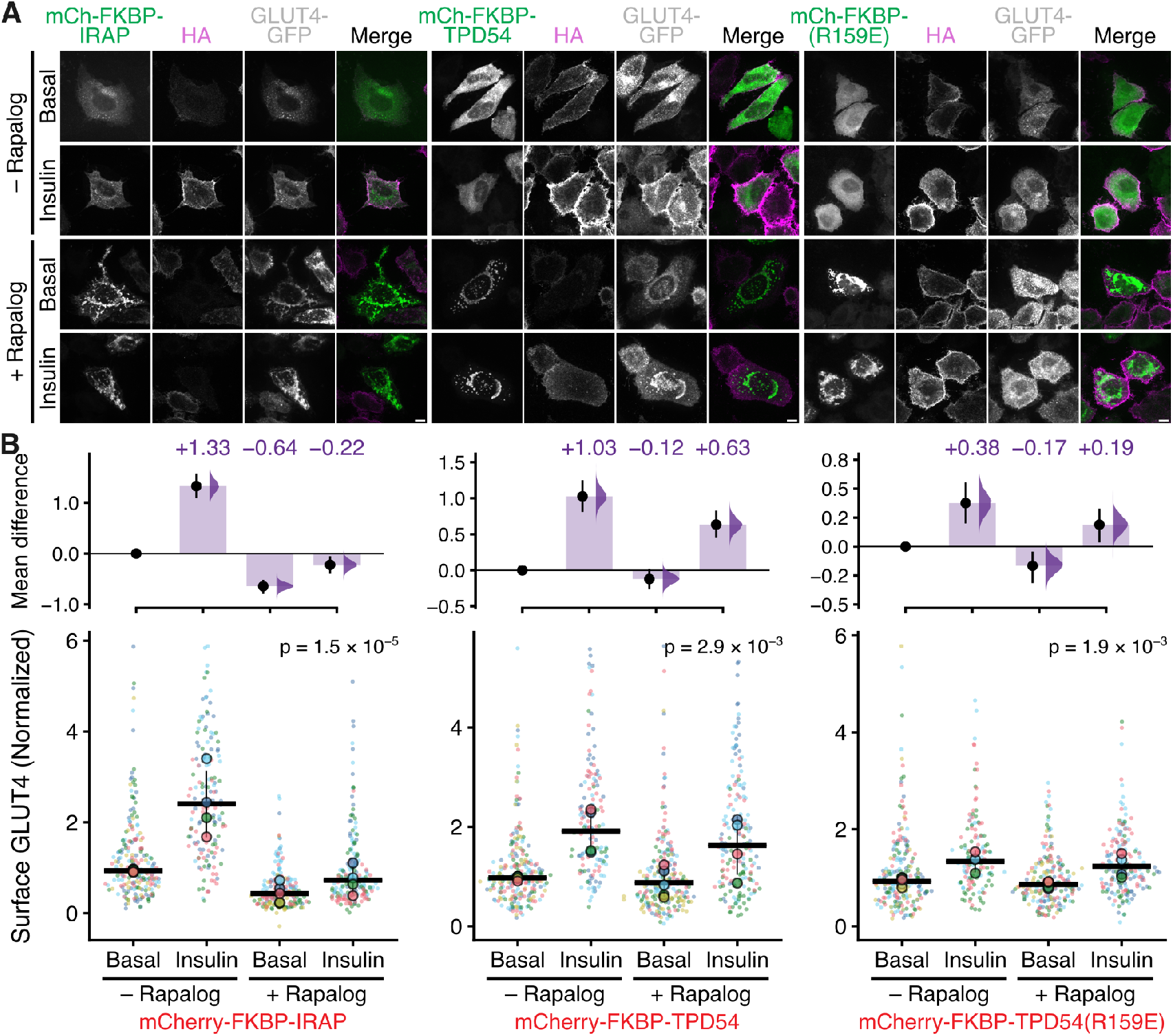
Capturing INVs at the mitochondria has only a minor effect on GLUT4’s insulin response. (**A**) Representative confocal images of HA-GLUT4-GFP HeLa cells, and have been co-transfected with dark Mitotrap (Mito–dCherry–FRB(T2098L)) and the mCherry-FKBP-tagged construct indicated. Cells were fixed and stained against surface GLUT4 using HA antibody. The merge image shows the mCherry-FKBP-tagged protein and HA staining). Images were taken with and without the addition of rapalog (5 µM, 10 min), followed by 20 min insulin stimulation (1 µg/mL) or without insulin (Basal). (**B**) Superplots showing the amount of GLUT4 at the cell surface as a proportion of total GLUT4, for each of the conditions. Data is normalized to the -Rapalog Basal condition for each mCherry-FKBP-tagged protein. Points, individual cells; colors, experimental repeat; circles, median for each repeat; bars, mean ± sd of the experimental medians. P-value, one-way ANOVA. Above: Cumming estimation plot to show effect sizes and bootstrap 95 % confidence interval.

We next performed vesicle capture of INVs using mCherry-FKBP-TPD54. Following insulin treatment, the redistribution of GLUT4 was not ablated but it was consistently lower than the redistribution seen following insulin treatment in cells with no capture (0.66 vs 1.15, or 57 % of the no-capture response) (Figure 8A,B). Indeed, the fraction of GLUT4 at the surface following capture was not significantly different from the basal condition. As a negative control, relocalization of the mCherry-FKBP-TPD54(R159E) mutant that is not vesicle associated had no effect on the insulin-induced redistribution of GLUT4 (Figure 8B).

These results suggest that GLUT4-containing INVs represent only a fraction of the insulin responsive pool of GLUT4 vesicles.

### GLUT4 INVs are important to the intracellular retention of GLUT4

GLUT4-containing INVs are a small fraction of the total INV population, so it is possible that the reason for incomplete inhibition of the insulin response was due to inefficient capture of GLUT4-INVs at the mitochondria. To address this possibility, we used RNAi to deplete TPD54 – which is a manipulation known to affect INV functions (Larocque et al., 2021; Fesenko et al., 2025) – to further test the role of INVs in insulinmediated GLUT4 redistribution. In HeLa cells stably expressing HA-GLUT-GFP, the insulin-induced redistribution of GLUT4 to the cell surface was robust with control siRNA (Figure 9A-C). Unexpectedly, we found that TPD54 knockdown caused an increase in the amount of GLUT4 at the surface under basal conditions, which was equivalent to 71 % of the insulin-stimulated redistribution in the control (Figure 9B). A small additional insulin-stimulated redistribution of GLUT4 was still observed in TPD54-depleted cells, but the increase was only 64 % of the control response, indicating that the GSV pool was smaller.

**Figure 9.**
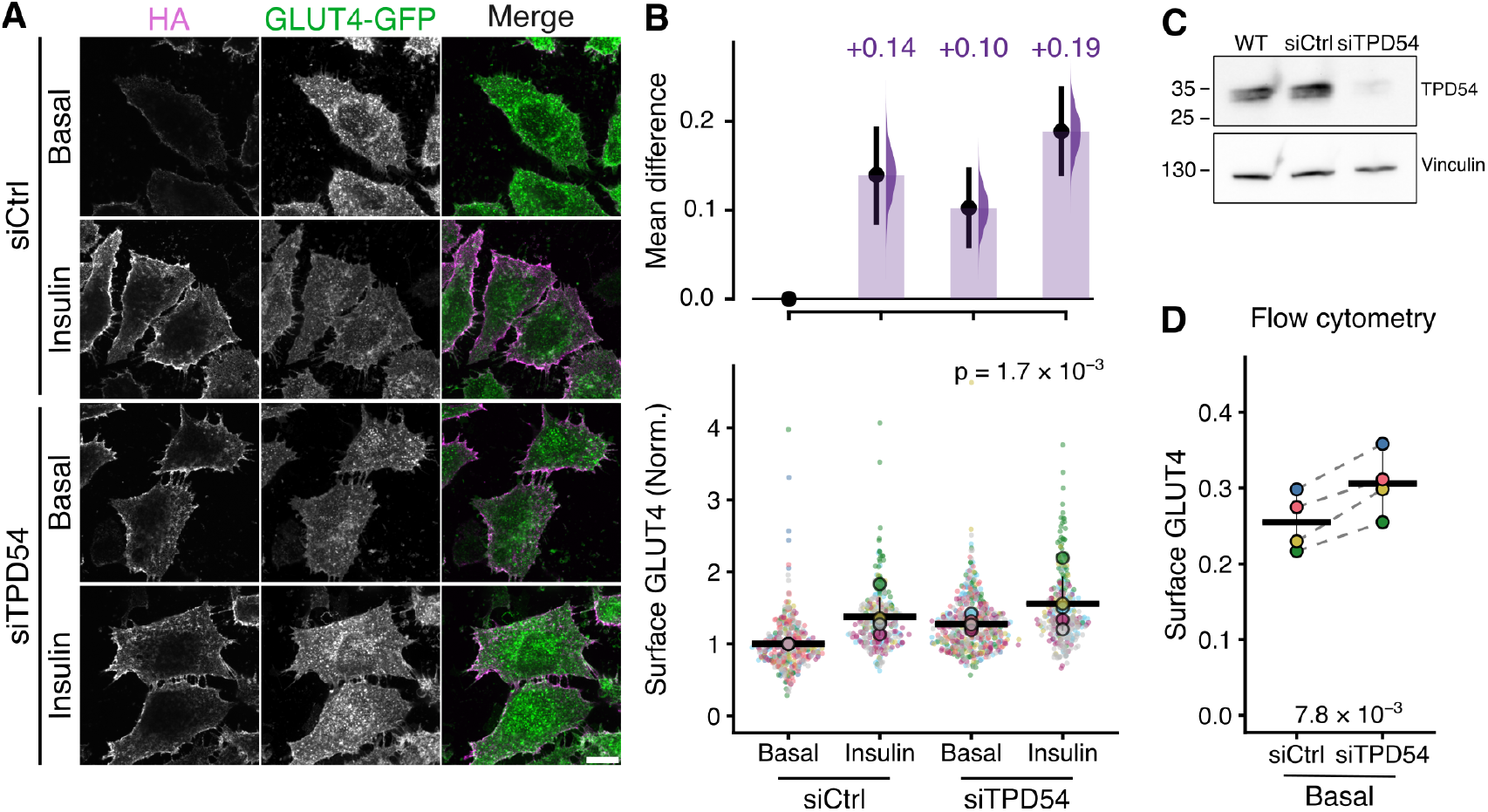
TPD54 knockdown increases surface GLUT4 in basal conditions. (**A**) Representative confocal images of HA-GLUT4-GFP HeLa cells following transfection with control (siCtrl) or TPD54 siRNA (siTPD54). Cells were serum starved for 2 h, followed by 20 min insulin stimulation (1 µg/mL) or not (Basal). Surface GLUT4 was visualized using anti-HA immunofluorescence (HA, magenta), GFP was used to assess total HA-GLUT4-GFP (green). (**B**) Superplot showing the amount of GLUT4 at the cell surface as a proportion of total GLUT4, for each of the conditions. Data are normalized to siCtrl Basal. Points, individual cells; colors, experimental repeat; circles, mean for each repeat; bars, mean ± sd of the experimental medians. P-value, one-way ANOVA. Above: Cumming estimation plot to show effect sizes and bootstrap 95 % confidence interval. (**C**) Western blot showing typical TPD54 levels following TPD54 knockdown. Loading control, vinculin. Uncropped blots are shown in Supplmentary Figure S2. (**D**) Flow cytometry measurement of the surface HA-GLUT4-GFP population, HA signal as a fraction of GFP fluorescence. The median is shown for each experimental repeat; bars, mean ± sd. P-value, Student’s paired t-test. 8.7 × 10^3^ to 1.8 × 10^4^ events were analyzed per condition per experiment.

Since the increase in GLUT4 surface levels in TPD54-depleted cells under basal conditions was modest, we used flow cytometry to measure the surface-to-total GLUT4 ratio. Again, we found a small but significant increase in this ratio when TPD54 was depleted (Figure 9D). This increase in surface GLUT4 could either be due to inhibition of endocytosis, or due to decreased sequestration of GLUT4 into the GSV pool, with more subsequent GLUT4 recycling to the plasma membrane. Since there is no known involvement of INVs in endocytosis (Larocque et al., 2020), we investigated this latter possibility.

Using expression of mCherry-Rab11(S25N) to inhibit recycling, we found that the increase in surface GLUT4 caused by TPD54 depletion was blocked by the dominant-negative construct when compared with expression of mCherry alone (Figure 10A). Expression of this construct in control siRNA-treated cells also decreased the surface GLUT4 population which highlights that GLUT4 is actively cycled between the PM and endosomes under basal conditions.

**Figure 10.**
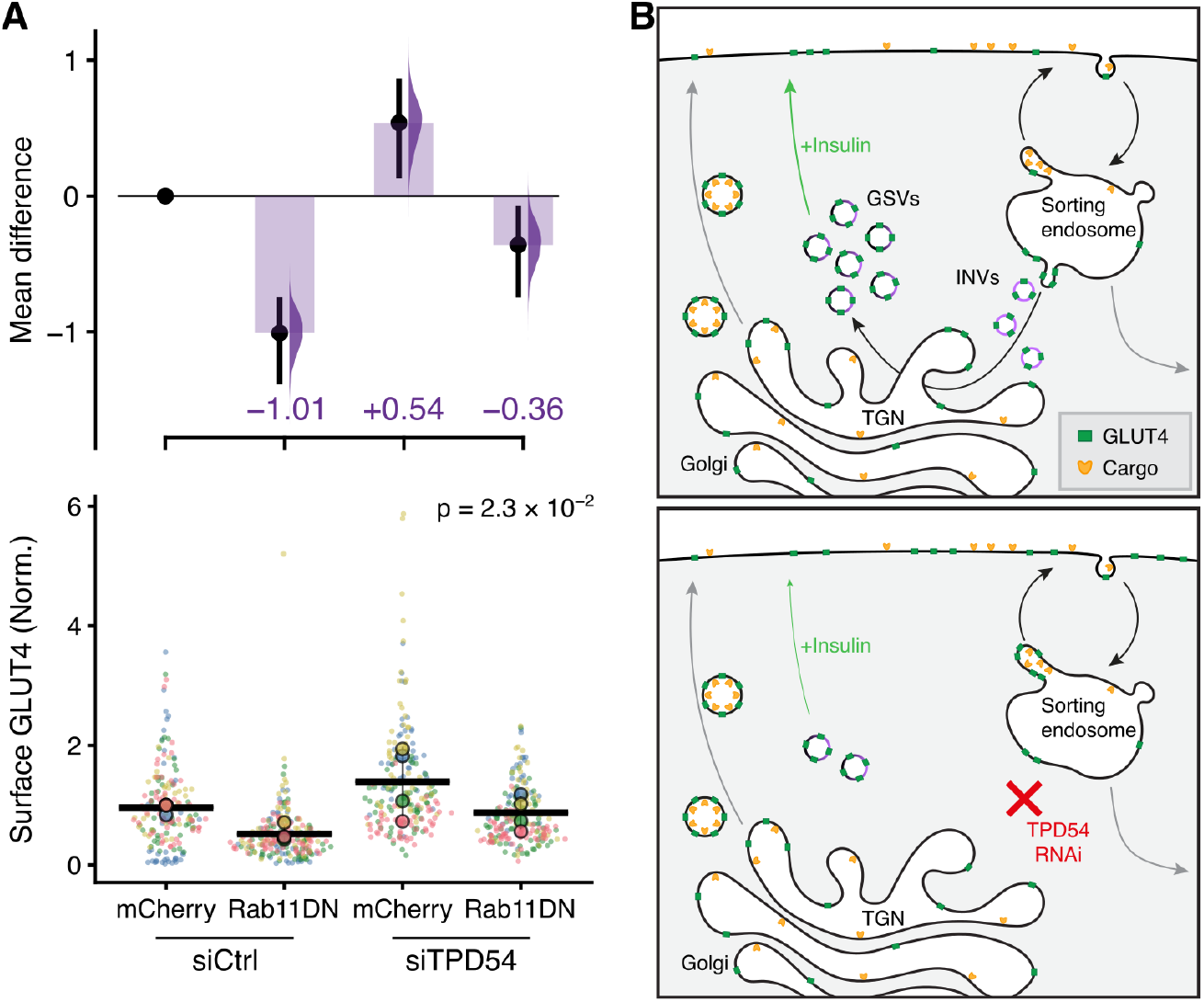
Blocking INV-mediated sequestration increases GLUT4 recycling to the plasma membrane. (**A**) Superplot showing the amount of GLUT4 at the cell surface as a proportion of total GLUT4, for each of the conditions. Cells were transfected with siCtrl or siTPD54 and expressed either mCherry or dominant-negative mCherry-Rab11(S25N) (Rab11DN). Data are normalized to siCtrl mCherry. Points, individual cells; colors, experimental repeat; circles, mean for each repeat; bars, mean ± sd of the experimental medians. P-value, one-way ANOVA. Above: Cumming estimation plot to show effect sizes and bootstrap 95 % confidence interval. (**B**) Schematic diagram of GLUT4 trafficking. Biosynthetic delivery and degradation of GLUT4 and other cargo (gray arrows) are balanced. Following endocytosis, GLUT4 is mainly sequestered from the sorting endosome to GSVs, whereas other cargo mainly recycles back to the cell surface. The GSV pool may be mobilized by insulin. We propose that GLUT4 sequestration is INV-mediated. When TPD54 is depleted, sequestration is impaired and GLUT4 is recycled with other cargo, increasing the surface population and reducing the GSV pool under basal conditions.

We propose a model where INVs are involved in a key retention step at the sorting endosome. GLUT4 is sorted away from the recycling pathway which would take it back to the plasma membrane, and instead INVs transport GLUT4 and associated cargo to form the insulin-responsive pool of vesicles (Figure 10B). In this model, disruption of INVs via TPD54 depletion, results in GLUT4 not being sorted into INVs and instead being constitutively recycled back to the plasma membrane. This would account for the increase in surface GLUT4 and the reduction in the GSV pool size observed in TPD54 knockdown cells.

## Discussion

Intracellular nanovesicles (INVs) are a large, molecularly diverse superfamily of vesicles operating on multiple membrane trafficking pathways in the cell. Here, we described how GLUT4, and other proteins associated with GLUT4 storage vesicles (GSVs), are carried by INVs. Functional assays indicated that INVs are primarily involved in GLUT4 retention – establishing and maintaining the insulin-responsive GSV pool – but that insulin-responsive GSVs themselves are not synonymous with INVs. These findings indicate a new role for INVs in sustaining the ability of the cell to mobilize GLUT4 in response to insulin.

Our previous proteomic and functional analysis of INVs had highlighted several molecular components commonly associated with GSVs (Larocque et al., 2020; Fesenko et al., 2025). Examples include Rab10 and Rab14, SCAMP1-3, R-SNAREs and SYNGR2 (Sano et al., 2011; Ishikura et al., 2007; Laurie et al., 1993; Volchuk et al., 1995; Kupriyanova and Kandror, 2000). INVs also carry several different transporters including GLUT1/SLC2A1, and are known to fuse with the plasma membrane (Fesenko et al., 2025; Sittewelle and Royle, 2024). It seemed likely therefore that GSVs were a subtype of INVs, in an analogous way to the ATG9A-flavor INVs that we had described previously (Fesenko et al., 2025). Consistent with this idea, our proteomic analysis of GLUT4-containing vesicles identified the INV marker TPD54, and a significant overlap with the INV proteome (Fesenko et al., 2025). However, some GSV components were not present in the overlap including IRAP/LNPEP, TBC1D4, syntaxins, LRP1 and GLUT4 itself. Their absence might be explained by the lack of GLUT4 expression in HeLa cells, where the INV proteome was determined; either via lower expression or reduced traffic (Kandror and Pilch, 2011). Using vesicle capture we confirmed that GLUT4, IRAP, cellugyrin, sortilin and VAMP2 are all present in INVs, and through a new single vesicle imaging method (co-occupancy), we determined that virtually all GLUT4 vesicle traffic is via INVs but that these GLUT4-flavor INVs are a small fraction of the total INV population. To directly test whether the insulin-responsive pool of GSVs were actually INVs we used mitochondrial vesicle capture to selectively remove defined populations of vesicles and test the effect on the insulin-induced redistribution of GLUT4 to the cell surface. While capture of IRAP vesicles abolished the insulin response, capture of INVs using TPD54 resulted in only a small impairment. This lack of phenocopying argues that the insulin-responsive GSVs cannot themselves be INVs – at most, INVs are ∼43 percent of this pool. Instead, depletion of TPD54, which interferes with INV function, resulted in deposition of GLUT4 to the cell surface and reduction of the size of the GSV pool. We can rule out the possibility that this is due to a block of GLUT4 endocytosis, because INVs are not involved in endocytosis of the transferrin receptor or α5β1 (Larocque et al., 2020, 2021). Instead this result points to a role for INVs in trafficking GLUT4 and related cargoes post-internalization to form the GSV pool.

There has long been evidence of two populations of GSVs, only one of which is insulin-responsive (Kupriyanova and Kandror, 2000; Kupriyanova et al., 2002; Kioumourtzoglou et al., 2015). The non-responsive pool likely represents precursor vesicles that are marked by the presence of cellugyrin/SYNGR2, SCAMPs and VAMP3 (Sevilla et al., 1997; Kupriyanova and Kandror, 2000; Jedrychowski et al., 2010). The fact that these proteins are all found in INVs, and that GLUT4-flavor INVs do not account for the entire insulin-responsive pool of GSVs, would suggest that GLUT4-flavor INVs are the precursor GSV population. Clearly, the delineation of vesicle pools is not binary. We found that sortilin is present in INVs, yet this protein is enriched in insulin-responsive GSVs (Kioumourtzoglou et al., 2015) and the capture of INVs did reduce the magnitude of GLUT4 redistribution in response to insulin. Any protein that is enriched in the insulin-responsive pool must traffic there via the precursor vesicles and similarly, capture of precursor vesicles is likely to impact the final insulin response to a varying extent. It is thought that all cells have the ability to sequester GLUT4 into an insulin-responsive pool of vesicles (Bryant and Gould, 2020), which indicates that this is achieved via the core membrane trafficking machinery. It is thought that GLUT4 retention occurs via IRAP-dependent sorting of GLUT4 at the endosome away from a ‘fast recycling pathway’ – by which proteins such as the transferrin receptor are recycled to the plasma membrane – into either a storage compartment such as the TGN, from which insulin-responsive GSVs are generated, or directly as the insulin-responsive GSVs themselves (Blot and McGraw, 2008; Govers et al., 2004; Jordens et al., 2010; Brumfield et al., 2021). Our work indicates that INV traffic represents the major route for the sequestration of GLUT4 as a key component of this core machinery, likely mediating traffic from the sorting endosome to the TGN as a precursor to generating the insulin-responsive GSVs.

Our study used mitochondrial vesicle capture to i) define molecular components of vesicles, ii) measure the size of molecularly-defined vesicles and iii) study the contribution of vesicles to a physiological response. The latter of these methods represents an exciting new way to study vesicle pools in cells and likely could be applied in other cell types and contexts. The capture of IRAP-positive vesicles at the mitochondria revealed their average diameter to be ∼44 nm, which is close to the 50 nm measured for GSVs following purification (Kandror et al., 1995), but slightly smaller than the 60– 100 nm measured by EM in rat adipocytes (Ramm et al., 2000). The size of these vesicles, like ATG9A-positive vesicles, is beyond the upper quartile of INV diameters that we measured. It is, however, still within the measured range for INVs, and could be easily accommodated within the INV population due to the fact that single vesicle imaging indicated that INV tracks outnumber GLUT4 tracks by greater than 9:1. Since co-occupancy analysis indicated that these vesicles are mainly INVs, it suggests an intriguing possibility that the size of INVs is related to how laden they are with cargo. Indeed, structural studies of ATG9A and subsequent molecular simulations show that its packing in a membrane causes deformation (Guardia et al., 2020). As an upper bound calculation, if the median INV size reflected no transmembrane cargo at all and the expansion to the median IRAP vesicle size was entirely due to GLUT4, this could be achieved with ∼240 copies of GLUT4. It is known that the magnitude of insulin-dependent redistribution of GLUT4 in rat adipocytes is much larger than in HeLa cells (Bryant and Gould, 2020) and this may simply reflect the amount of GLUT4 packing into the GSVs, as suggested by the smaller diameters measured here (∼44 nm) versus in rat adipocytes 50–100 nm. The direct visualization of sizes corresponded well to our single vesicle imaging of diffusional mobility. These single vesicle approaches also revealed a decrease in co-occupancy between GLUT4 and TPD54 when cells were stimulated with insulin, an observation which suggests that GLUT4 localization shifts from the precursor pool, which supports our conclusion that GLUT4-flavor INVs are largely separate from the insulin-responsive GSVs. Together vesicle capture, single vesicle imaging of co-occupancy and mobility represent powerful new methods to apply to the study of GLUT4 trafficking and of related questions in membrane traffic by delineating defined vesicle pools in live cells.

## Methods

### Molecular biology

The following plasmids were available from previous work: mCherry-FKBP-tagged or GFP-tagged TPD54 WT and R159E mutant, GFP-VAMP2, and mCherry-FKBP (Larocque et al., 2020, 2021); StayGold-TPD54 (Sittewelle and Royle, 2024); blue MitoTrap (pMito-EBFP2-FRB[T2098L]), dark MitoTrap (pMito-dCherry-FRB[T2098L]), and ATG9A-FKBP-mCherry (Fesenko et al., 2025); green MitoTrap (pMito-GFP-FRB) (Clarke and Royle, 2018).

SYNGR2 and Sortilin tagged with monomeric GFP were made by DNA synthesis. HA-GLUT4-GFP and IRAP-pHluorin were kind gifts from Karin Stenkula (Lund). mCherry-FKBP-IRAP was made by PCR of IRAP from IRAP-pHluorin to add MfeI and EcoRI sites and reintroduced the stop codon before cloning into a vector containing mCherry-FKBP. ATG9A-StayGold, ATG9A-mScarlet-I3, HA-GLUT4-StayGold, and HA-GLUT4-mScarlet-I3 were made by amplification of StayGold or mScarlet-I3, to add KpnI and MfeI sites and cloning in place of mCherry in ATG9A-mCherry or GFP in HA-GLUT4-GFP. mScarlet-I3-TPD54 was made by PCR of mScarlet-I3 to add AgeI and BsrGI and replace GFP in GFP-TPD54. To make mCherry-Rab11a S25N, GFP-Rab11a S25N was made by site directed mutagenesis from GFP-Rab11a (gift from Francis Barr, Oxford), and the fluorescent protein was swapped to mCherry by digesting pmCherry-C1 with AgeI and BsrGI.

### Cell culture

Wild-type HeLa cells (HPA/ECACC 93021013), GFP-TPD54 knock-in (clone 35) HeLa cells (Larocque et al., 2020) or HeLa cells stably expressing HA-GLUT4-GFP (gift from Gwyn Gould, Strathclyde) were maintained in DMEM with GlutaMAX and 25 mM HEPES (Thermo Fisher, 32430100) supplemented with 10 % FBS, and 100 U mL^−1^ penicillin/streptomycin. All cells were kept in a humidified incubator at 37 °C and 5 % CO_2_; and were routinely tested for mycoplasma contamination by PCR.

For transient transfection, 2 × 10^5^ cells were plated onto either cover slips or 35 mm glass bottom dishes (WPI – FD35-100). Plasmids were transfected using Gene-juice and cells analyzed 36–48 h after transfection. For depletion of TPD54, siTPD54 (GUCCUACCUGUUACG-CAAU) or siGL2 (CGUACGCGGAAUACUUCGA) as a control were used (Larocque et al., 2020). Transfection of siRNA was by Lipofectamine RNAiMax according to the manufacturer’s instructions, with cells typically analyzed 48 h post-transfection.

### Proteomic analysis of HA-GLUT4-GFP vesicles

HA-GLUT4-GFP or WT HeLa cells (14 × 10^6^) were seeded onto 24.5 × 24.5 cm plates (Corning, 431110). Typically one plate per condition, per replicate was set up and left to grow until 80–100 % confluent. For harvest, plates were placed on ice, cells were washed three times with 10 mL ice cold PBS and scraped into 5 mL PBS using a cell lifter. Cells were pelleted by centrifugation at 4200 rpm for 2 min at 4 °C and then lysed via resuspension of the cell pellet in 1 mL vesicle release buffer (20 mM Tris-Cl, 300 mM NaCl, 100 µg mL^−1^ digitonin, 0.2 mM PMSF, with cOmplete EDTA free proteinase inhibitor (Roche, 4693132001), pH 7.5) in a pre-cooled microcentrifuge tube. Cells were lysed on ice for 30 min, with gentle pipetting every 10 min using a 1 mL tip. During lysis, GFP-Trap Agarose beads (chromotek, gta-20), 50 µL per reaction, were prepared by washing with 500 mL INV release buffer with spins at 2500 *g* for 2 min at 4 °C). Lysates were then centrifuged at 20 000 *g* for 15 min at 4 °C, 50 µL supernatant was reserved for analysis and the remainder was incubated with GFP-Trap beads (50 µL per reaction) in a pre-cooled microcentrifuge tube and incubated with end-over-end rotation at 4 °C for 1 h. The bead-lysate mixture was centrifuged at 2500 *g* for 2 min at 4 °C to pellet the beads. The supernatant was retained for analysis and the vesicles on beads were washed three times with vesicle release buffer (without digitonin). Captured material was eluted by resuspending beads in 50 µL 2× Laemmli buffer (Alfa Aesar, J60015.AD) and heating to 95 °C for 5 min.

Samples were loaded onto a 4–15 % precast polyacrylamide gel (Bio-Rad, #4561084) and run at constant voltage until all proteins had migrated into the resolving gel. The gel was stained with Coomassie Blue stain (50% Methanol, 10% Acetic acid, 0.1% Coomassie Brilliant Blue R-250) for 30 min at room temperature (RT) with gentle agitation, and de-stained overnight (15% Methanol, 15% Acetic acid). Individual lanes were excised, diced (2–4 mm), and transferred to 1.5 mL microcentrifuge tubes per lane. Samples were then prepared for mass spectrometry as in (Fesenko et al., 2025)

LC-MS/MS raw data files were processed using MaxQuant (v2.0.1) (Max Planck Institute of Biochemistry). The peptide lists were searched against the reviewed UniProt human proteome database (retrieved May 2021) supplemented with the sequence of HA-GLUT4-GFP and the MaxQuant common contaminant database. Peptide abundance was quantified using the MaxQuant Label-Free Quantification (LFQ) algorithm. Enzyme specificity for trypsin was selected with the allowance of up to two missed cleavages. All searches were performed with cysteine carbamidomethylation set as the fixed modification and oxidation of methionine and acetylation of the protein N-terminus set as the variable modifications. The initial precursor mass deviation was set as 20 ppm and the fragment mass deviation set as 20 ppm. For label-free quantification, we set a minimum ratio count of 2 with 3 minimum and 6 average comparisons.

### Cell treatments

To capture vesicles at the mitochondria, an inducible heterodimerization approach was used as described previously, with minor modifications (Larocque et al., 2020). Briefly, cells expressing FKBP-tagged proteins and FRB T2098L-MitoTrap constructs were treated with rapalog AP21967 (A/C Heterodimerizer, TaKaRa Bio, 635056) at a final concentration of 5 µM in media.

For live cell imaging, cells were in Leibovitz L-15 media supplemented with 10 % FBS immediately prior to imaging and rapalog application was done using a 1:5 addition of a 5X stock in L-15 media with FBS. Insulin stimulation was done using a 1:2 addition of L-15 media containing 2 µg/mL insulin.

For vesicle capture with subsequent insulin stimulation, cells were serum-starved using DMEM with GlutaMAX and 25 mM HEPES (Thermo Fisher, 32430100) without any supplements. After 90 min, rapalog was applied via complete exchange of media for 10 min. Following this, 1 µg/mL insulin (± rapalog) was applied by complete exchange of media for 10 min. Cells were then put on ice prior to fixation and antibody staining.

### Immunofluorescence

Cells on glass cover slips were fixed using 3 % PFA/4 % sucrose in PBS for 15 min. Cells were washed twice with PBS, before 45–60 min blocking (3 % BSA, 5 % goat serum in 1xPBS). Antibody dilutions were prepared in blocking solution. After blocking, cells were incubated for 1 h with primary antibody, washed with PBS (3 washes, 5 min each), 1 h secondary antibody incubation, 5 min each PBS wash (3 washes), mounting with Vectashield vibrance. Primary antibody: anti HA.11 [6E2], mouse (Biolegend, 901513); Secondary antibody: Alexa Fluor647 conjugated goat anti-mouse, highly cross-adsorbed (Thermo-Fisher).

For flow cytometry, cells were plated into a 6-well plate and a blocking solution of 2 % BSA was used. Following antibody staining and washing (as described), cells were lifted from the plate using a cell lifter, and spun at 600 *g* for 8 min. The supernatant was then removed and replaced with 0.5 mL of blocking solution. Samples were then analyzed using the BD LSRFortessa using 488 nm and 640 nm lasers with Green 530/30 and FRed 670/30 emission filters. 10 × 10^4^ cells were captured per condition per replicate.

### Western blotting

Cell samples were resolved on a precast 4–15 % polyacrylamide gel (Bio-Rad) and transferred to nitrocellulose using a iBlot2 Dry Blotting System (Bio-Rad). Following blocking in 5 % w/v non-fat milk (Merck, 70166) in TBST buffer (20 mM Tris, 150 mM NaCl, 0.1 % v/v Tween-20, pH 7.6), membranes were incubated with primary antibodies: rabbit polyclonal anti-vinculin (Sigma Aldrich, V4139) or rabbit polyclonal anti-TPD54 (Dundee Cell Products) at 1:1000 in 2.5 % milk-TBST for 2 h at RT or at 4 °C overnight with agitation. After three washes in TBST, secondary antibodies were applied at 1:2000 in 5 % milk TBST for 1 h at RT with agitation: HRP-conjugated sheep anti-mouse IgG (Cytiva, NXA931V) or mouse anti-rabbit IgG (Santa Cruz Biotechnology, sc-2357). Blots were imaged using Amersham ECL Prime Western Blotting Detection Reagent (Cytiva, RPN2236) on a ChemiDoc MP (Bio-RAD) digital imaging system.

### Microscopy

All images were captured using a Nikon CSU-W1 spinning disc confocal system with SoRa upgrade (Yokogawa) with a Nikon, 100×, 1.49 NA, oil, CFI SR HP Apo TIRF with optional 2.8× intermediate magnification and a 95B Prime camera (Photometrics). The system has a CSU-W1 (Yokogawa) spinning disk unit with 50 µm and SoRa disks (SoRa disk used), Nikon Perfect Focus autofocus, Okolab microscope incubator, Nikon motorized xy stage and Nikon 200 µm z-piezo. Excitation was via 405 nm, 488 nm, 561 nm and 638 nm lasers with 405/488/561/640 nm dichroic and Blue, 446/60; Green, 525/50; Red, 600/52; FRed, 708/75 emission filters. For vesicle tracking, a two camera setup was used for simultaneous capture of both channels at a high frame rate. All other imaging was done using single camera, sequential capture. Acquisition and image capture was via NiS Elements software (Nikon). All microscopy data was stored via automated nightly upload to an OMERO database in the native file format (nd2).

### Electron Microscopy

HeLa cells were seeded onto gridded dishes (P35G-1.5-14-CGRD, MatTek) and transfected with MitoTrap (pMito-GFP-FRB) and either mCherry-FKBP, mCherry-FKBP-TPD54, ATG9A-FKBP-mCherry or mCherry-FKBP-IRAP. 24 h post-transfection, cells of interest were selected by light microscopy with a SoRa confocal microscope and capture of the FKBP-tagged protein at the mitochondria was done by adding 200 nM rapamycin. Vesicle capture was achieved after 15 min of rapamycin treatment and the cells were then fixed for 24 h at 4 °C with PHEM buffer (60 mM PIPES, 25 mM HEPES, 10 mM EGTA, 2 mM MgCl_2_, pH 6.9) containing 1 % glutaraldehyde. The aldehydes were then quenched with 100 mM glycine in HEPES 0.2 M for 10 min and washed with water. The cells were stained for 1 h with 1 % OsO_4_ and 1.5 % potassium ferricyanide at room temperature, washed with water and stained further with 1 % uranyl acetate (UA) at 4 °C overnight. The cells were washed again with water and gradually dehydrated with 10 min incubations at RT with ethanol (EtOH) at different concentrations (50, 70, 80, 90, 96 and 100 %) (an additional staining step of 1 % UA in 70 % EtOH for 1 h and washes in 70 % EtOH were included between the incubations in 70 and 80 % EtOH) and embedded in epon resin (EMS). Ultrathin sections were cut at a 50 nm thickness on a EM UC7 (Leica Microsystems), collected on formvar-coated grids and stained for 1 h in 4 % UA in 50 % EtOH and 30 s in Reynold’s lead citrate. The cells of interest were located and imaged using a JEOL2100Plus with a LaB6 200 kV filament fitted with a Gatan OneView IS camera.

### Data analysis

Data from MaxQuant was processed using VolcanoPlot in Igor Pro 9 (Royle, 2024). Cut-offs of two-fold enrichment and *p* < 0.05 were used to define proteins enriched in HA-GLUT4-GFP vesicles. Comparison with the INV proteome was done in R using custom-written routines (see below).

For analysis of relocalization–co-relocation experiments, three-channel images (FKBP protein, POI, Mitotrap) were first registered with NanoJ using TetraSpeck bead images. The mitochondrial channel was segmented using a Labkit classifier trained on MitoTrap images (Arzt et al., 2022). Fluorescence measured in the mitochondrial region post-treatment was divided by the pre-treatment value (F_post_/F_pre_), for each cell.

For vesicle co-occupancy analysis, simultaneous two-channel capture movies at 0.09 s per frame were analyzed using TrackMate (Ershov et al., 2022). An automated procedure tracked spots in each channel in turn, within an ROI that contained the cell of interest. Since tracking relies on unambiguous detection and frame-to-frame linkage, not all vesicles are tracked. This means that the coincidence of tracks cannot be used for co-occupancy analysis because they are incomplete. However, the tracks can be used to define vesicle signal from background in each channel and therefore to determine co-occupancy in unambiguous tracks in the other channel. Accordingly, the fluorescence intensities of tracks recorded in each primary channel were used to determine the mean ± sd of the signal attributable to vesicles in that channel. This information was used to set a threshold to determine if the corresponding secondary channel was positive (colocalized) or not for that signal, in that frame, when the other (primary) channel was tracked. The threshold was set as 0 < *σ* ≤ 2 below the mean, the magnitude of which was determined by randomizing the pixels in the secondary channel and substituting the mean fluorescence for the actual secondary signal assessing colocalization; and then determining the largest *σ* value that yielded < 0.05 co-occupancy, i.e. an error rate of 5 %. A minimum track length of 10 frames was used and colocalization of 80 % of frames was used to define a co-occupied track. This approach i) accounted for photobleaching, 2) was independent of fluorescence intensity differences between constructs or between cells, iii) was more robust than other colocalization measures since it measured the persistence of signals over time, and iv) was applied at the level of single vesicles. The approach was validated using synthetic data (Supplementary Figure S1). Briefly, movies with 400 “vesicles” diffusing in 2D with a diffusion coefficient of 0.3 µm^2^ s^−1^ were generated using the same imaging parameters as the live cell imaging. Background and vesicle signals were comparable to the experimental data. For a co-occupancy of 0.2, 80 vesicles were co-occupied while 160 were channel 1-positive and 160 were channel 2-positive. This results in a per channel co-occupancy of 0.3 (80 out of 240), rather than 0.2.

Analysis of vesicle mobility was done using an automated TrackMate (Ershov et al., 2022) procedure and subsequent processing of TrackMate XML files using TrackMateR (Royle, 2022) v0.3.12.

For vesicle capture analysis, electron micrographs were manually segmented in IMOD. IMOD models were converted using model2point and imported into R. Area and perimeter were found and used to calculate circularity according to

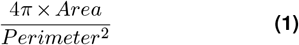

and a cut off of > 0.7 used for vesicles. For each vesicle pointset, the mean polar radius relative to the centroid was used to calculate the diameter. PCA was used to rotate the pointset and align vesicles along their major axis, and a random selection of 150 profiles taken for display.

All image analysis outputs from Fiji were read into R, analyzed and plotted using custom-written scripts. Superplots were generated in R using SuperPlotR v0.0.8 (Royle, 2025) and effect size plots were made using dabestr (Ho et al., 2019).

## Data and software availability

All code used in the manuscript is available at https://github.com/roylelab/p070p039 and archived at https://doi.org/10.5281/zenodo.18661081. Data are either archived with the code or at https://doi.org/10.5281/zenodo.18660082

## ACKNOWLEDGEMENTS

We thank Gwyn Gould for the HA-GLUT4-GFP cell line, and Francis Barr and Karin Stenkula for plasmids. We acknowledge the invaluable support of our core facilities: Warwick Proteomics RTP, Advanced Bioimaging RTP (EM), Flow Cytometry SRL, and Computing and Advanced Microscopy Unit (CAMDU). We are grateful to all members of the Royle lab for feedback and critical discussion. The work was supported by grants from UKRI-BBSRC (BB/V003062/1) and Human Frontier Science Program (HFSP RGP25/2022). For the purpose of open access, the authors have applied a Creative Commons Attribution (CC BY) license to any Author Accepted Manuscript version arising

## AUTHOR CONTRIBUTIONS

Investigation: EC and GL; Software: EC and SJR; Formal analysis: EC and SJR; Visualization: EC, GL and SJR; Funding acquisition: SJR; Writing (original draft): EC; Writing (reviewing and editing): EC, GL and SJR.

## COMPETING FINANCIAL INTERESTS

The authors declare no conflict of interest.

## Supplementary Information

**Figure S1.**
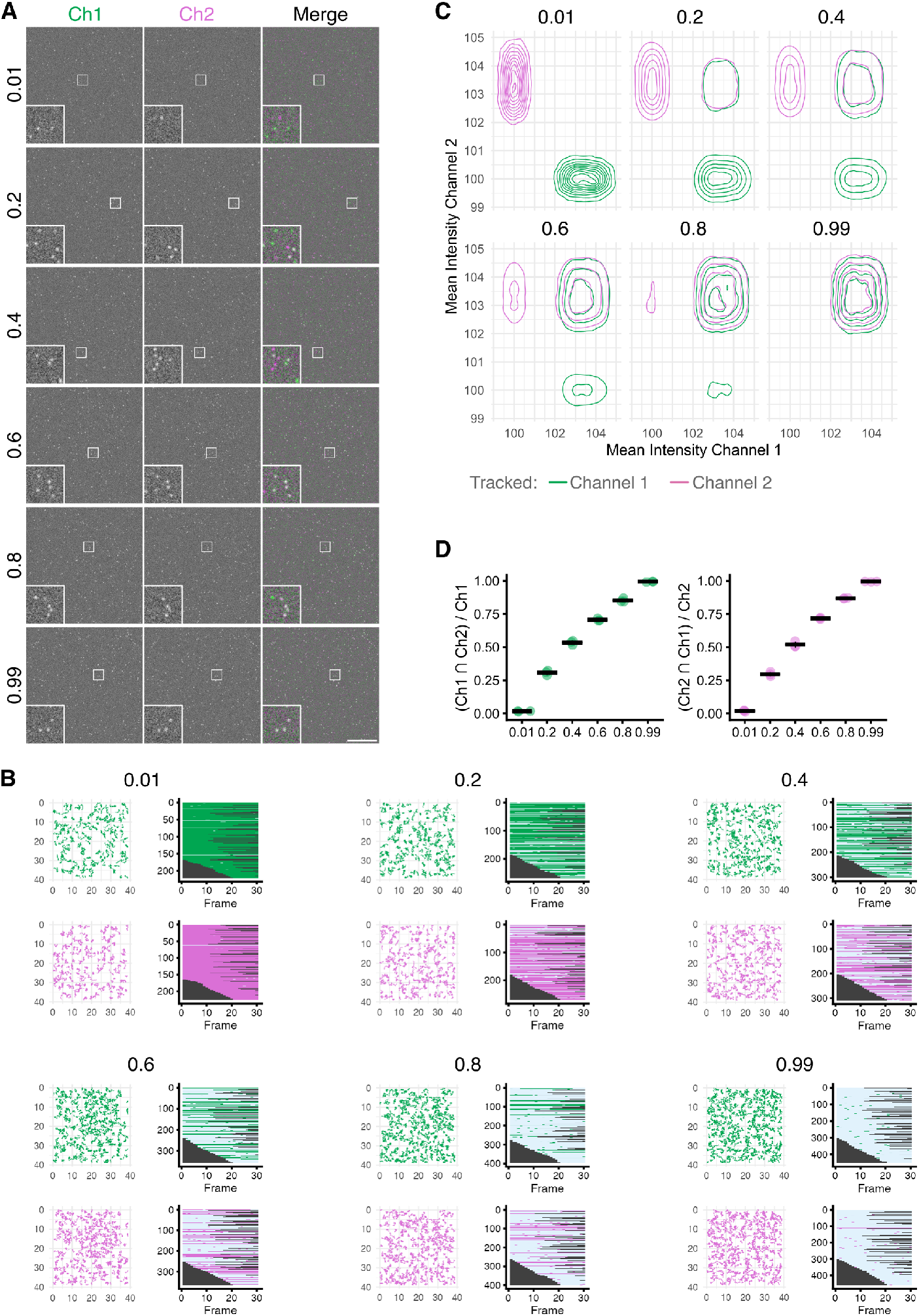
Validation of single vesicle co-occupancy determination. (**A**) Stills from synthetic movies generated with the indicated fraction of co-occupancy between channel 1 (Ch1) and channel 2 (Ch2). (**B**) Tracking results from a single movie at the indicated fraction of co-occupancy. Using the workflow, each event is assessed for colocalization at each frame and a level of > 80 % of frames with colocalization (light blue) is required to designate the vesicle as co-occupied. (**C**) The intensities of tracked vesicles and their corresponding locations is shown for all movies in the dataset, (**D**) Measured per channel co-occupancy versus population co-occupancy, determined by the workflow. Expected values are 0.02, 0.3, 0.57, 0.75, 0.89, and 0.99. Spots are individual movies, bars indicate the mean ± sd.

**Figure S2.**
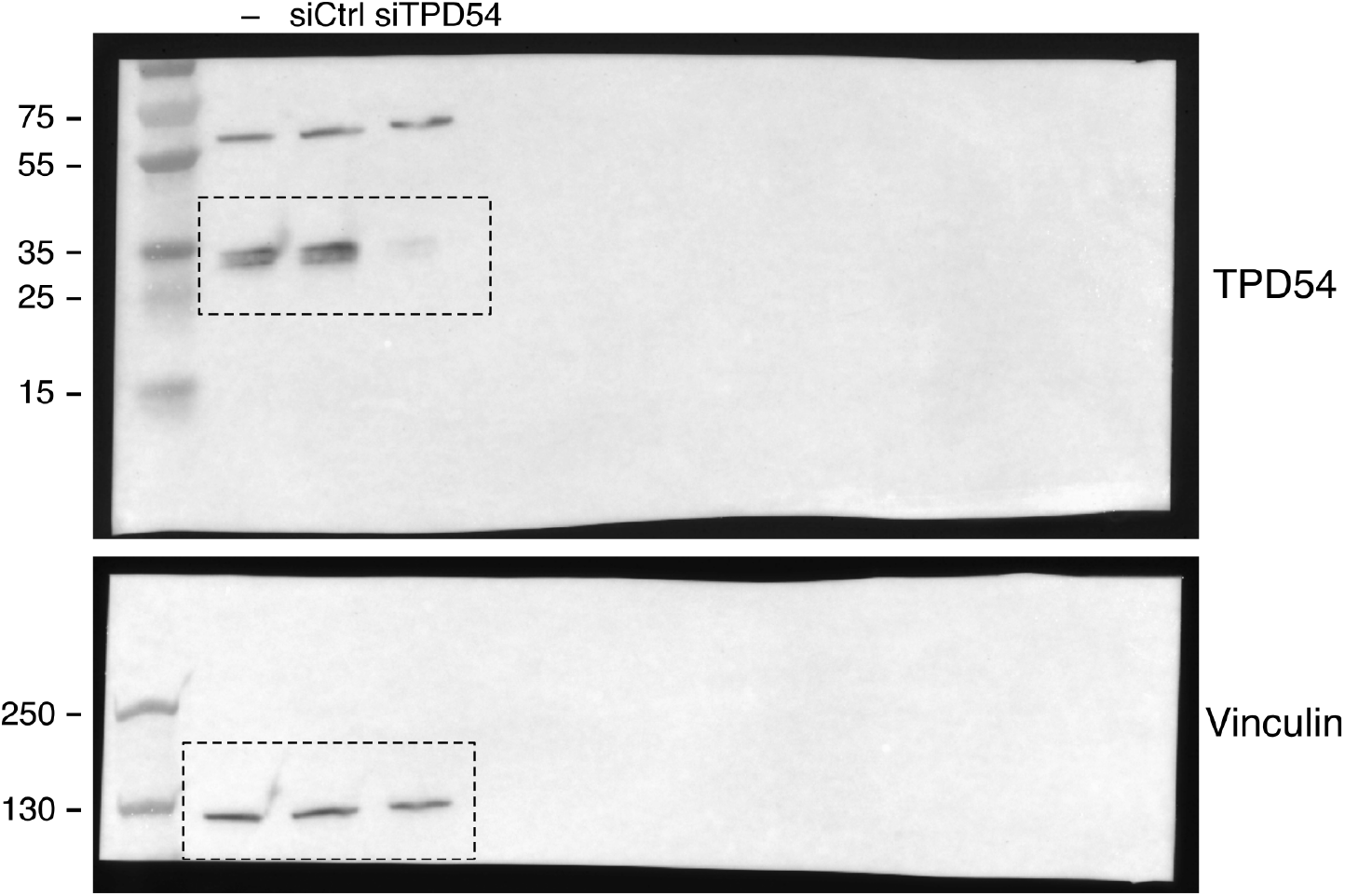
Uncropped western blots from Figure 9

## Supplementary Videos

**Figure SV1.**
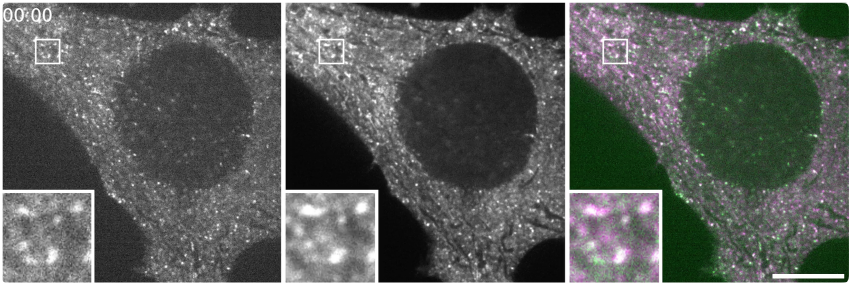
Vesicle co-occupancy: TPD54 with TPD54. HeLa cell co-expressing StayGold-TPD54 (left, green), mScarlet-I3-TPD54 (middle, magenta), merge (right). Zooms, 4X; Playback, 10 fps; Time, mm:ss; Scale bar, 10 µm.

**Figure SV2.**
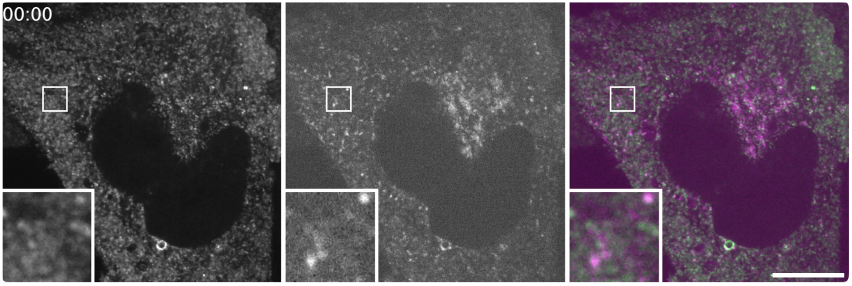
Vesicle co-occupancy: TPD54 with GLUT4. HeLa cell co-expressing StayGold-TPD54 (left, green), GLUT4-mScarlet-I3 (middle, magenta), merge (right). Zooms, 4X; Playback, 10 fps; Time, mm:ss; Scale bar, 10 µm.

**Figure SV3.**
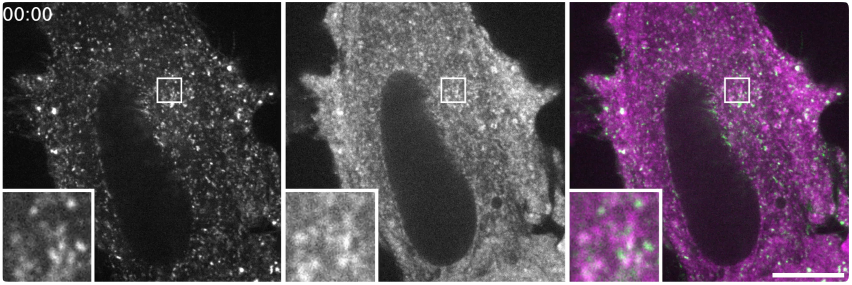
Vesicle co-occupancy: GLUT4 with TPD54. HeLa cell co-expressing GLUT4-StayGold (left, green), mScarlet-I3-TPD54 (middle, magenta), merge (right). A second playback illustrates the individual vesicle tracking results. Zooms, 4X; Playback, 10 fps; Time, mm:ss; Scale bar, 10 µm.

**Figure SV4.**
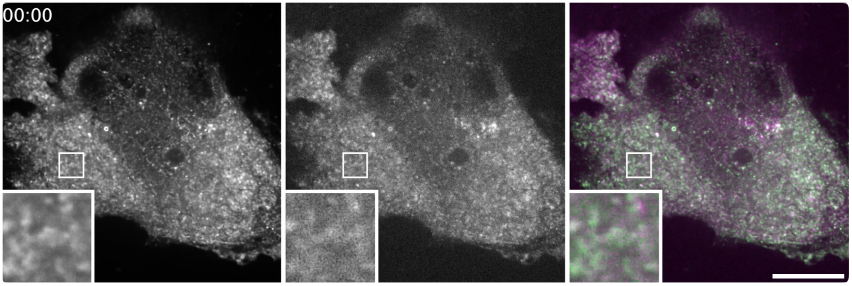
Vesicle co-occupancy: TPD54 with ATG9A. HeLa cell co-expressing StayGold-TPD54 (left, green), ATG9A-mScarlet-I3 (middle, magenta), merge (right). Zooms, 4X; Playback, 10 fps; Time, mm:ss; Scale bar, 10 µm.

**Figure SV5.**
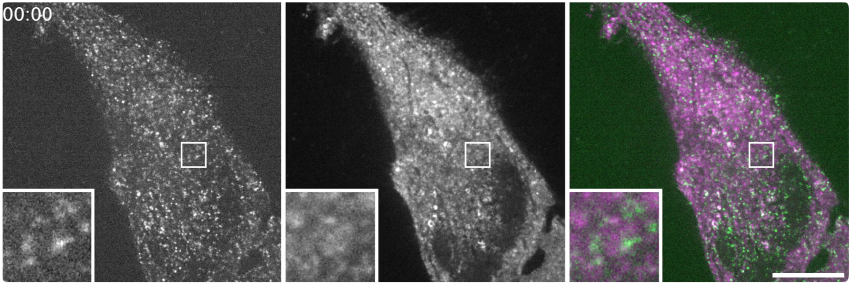
Vesicle co-occupancy: ATG9A with TPD54. HeLa cell co-expressing ATG9A-StayGold (left, green), mScarlet-I3-TPD54 (middle, magenta), merge (right). Zooms, 4X; Playback, 10 fps; Time, mm:ss; Scale bar, 10 µm.

## Supplementary Tables

**Table S1.**
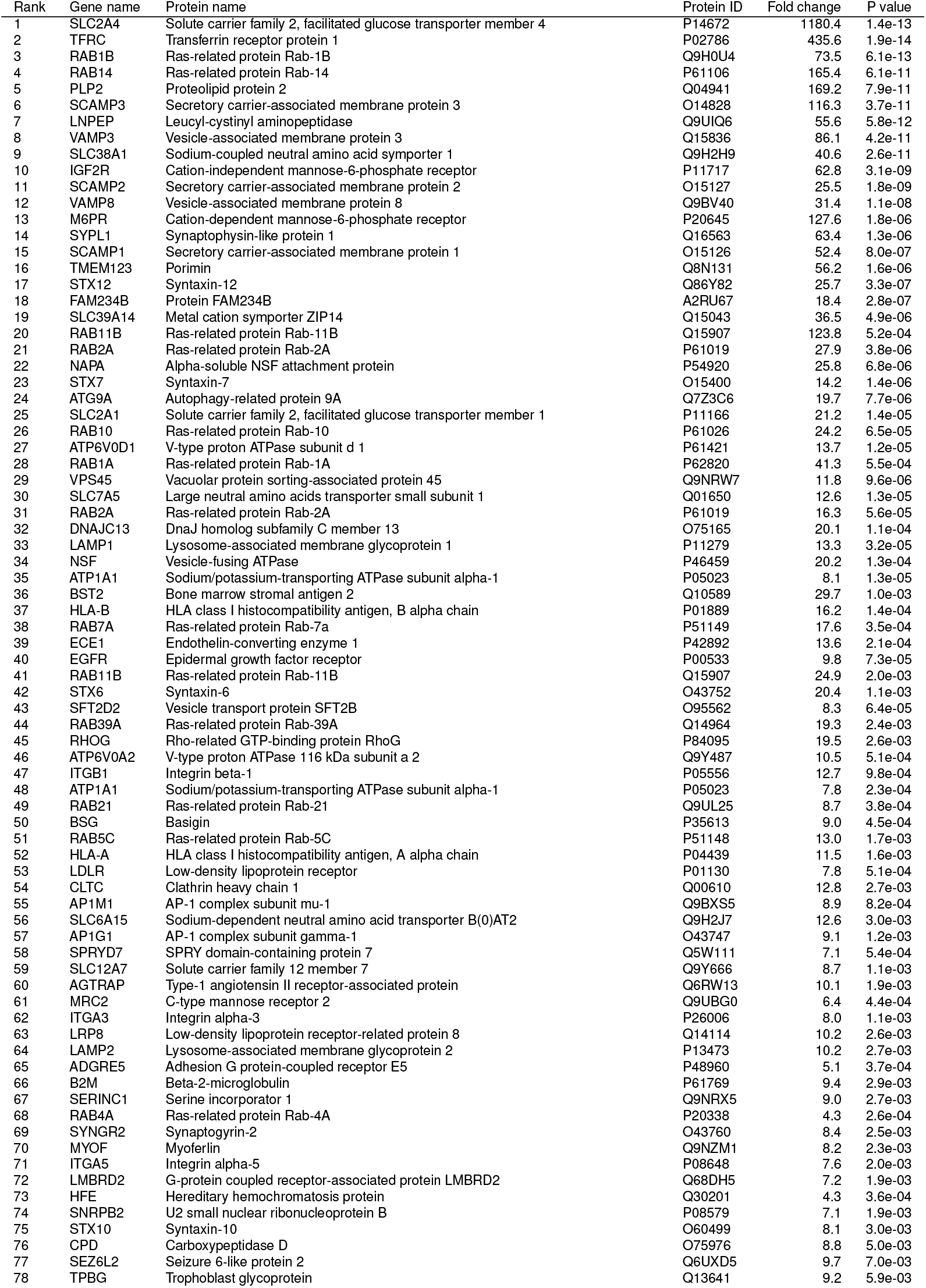
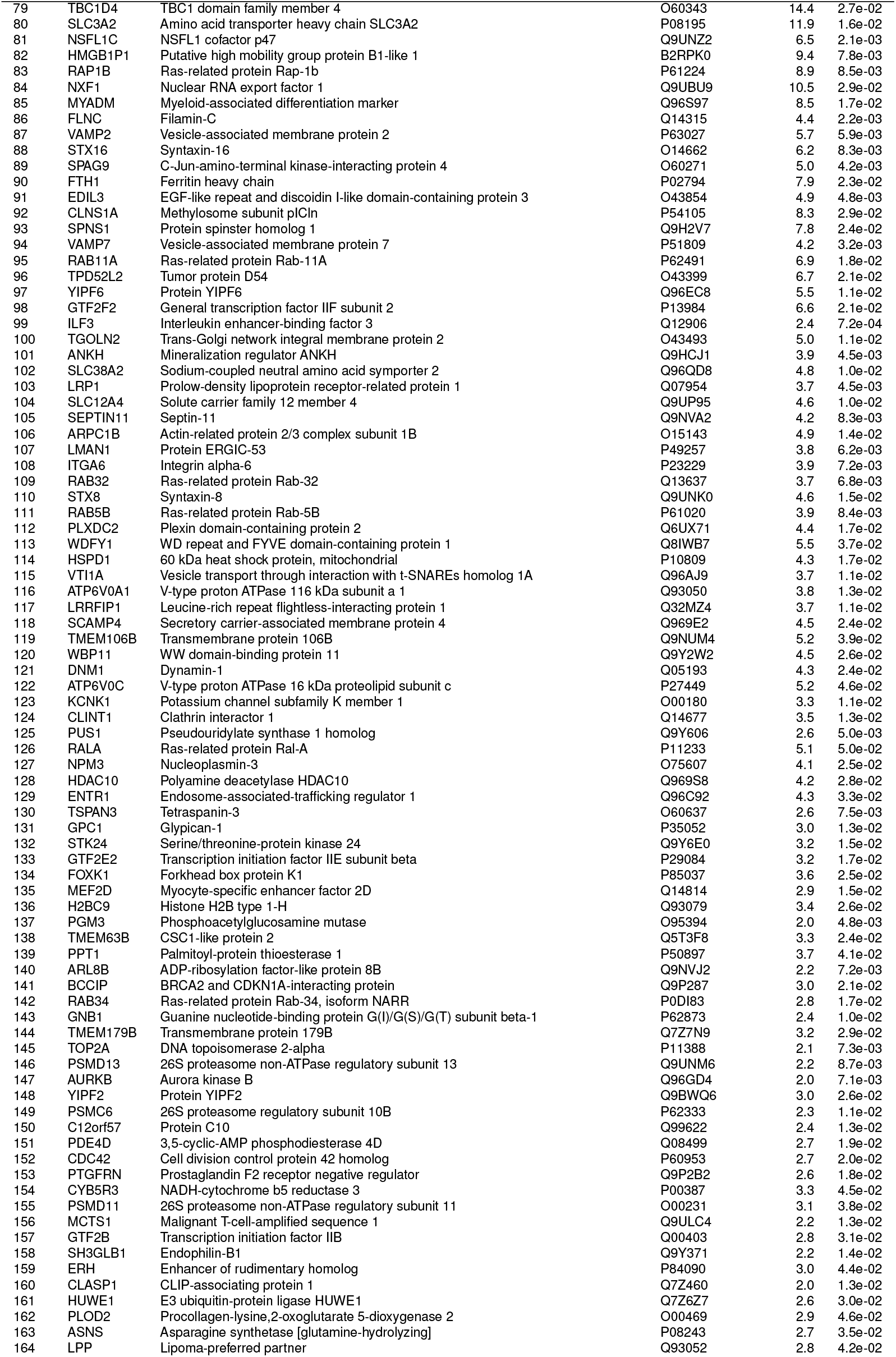
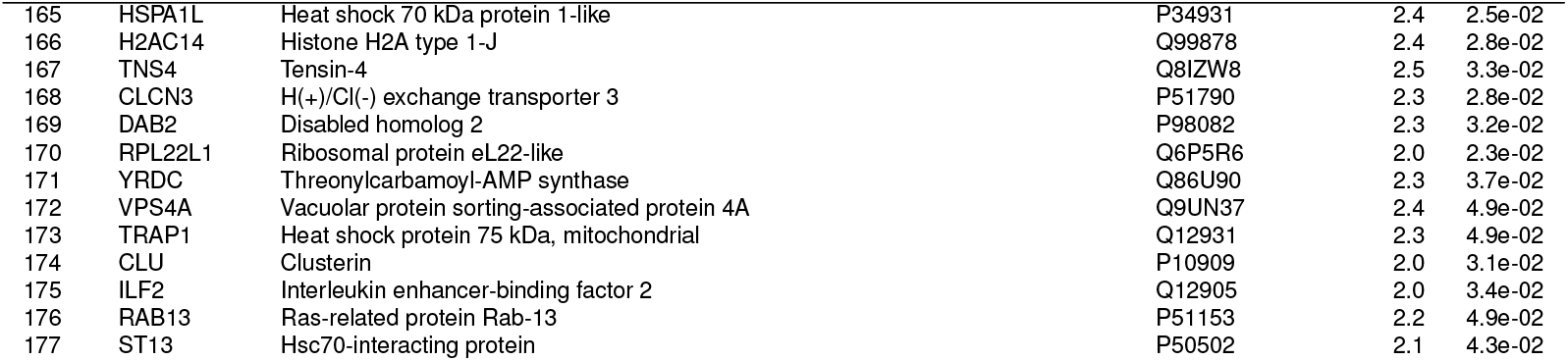
Significant hits from HA-GLUT-GFP proteomics. A list of the proteins ranked by their fold enrichment over control. 177 proteins that had a fold change of > 2 and p < 0.05 are included. See Figure 1. For entries with more than one identifier, only the first is shown for clarity.

## Notes

https://doi.org/10.5281/zenodo.18661081

https://doi.org/10.5281/zenodo.18660082

### Competing Interest Statement

The authors have declared no competing interest.

